# Tree-Cover Gradients Modulate the Taxonomic and Functional Diversity of Birds in Amazonian Cattle-Ranching Landscapes

**DOI:** 10.1101/2025.11.07.687096

**Authors:** Alexander Velasquez-Valencia, Jenniffer Tatiana Díaz-Cháux, María Argenis Bonilla-Gómez, Alejandro Navarro-Morales, Lina Paola Giraldo, Fernando Casanoves

## Abstract

Cattle-ranching expansion is the major anthropogenic driver transforming forests in the Colombian Amazon. Landscapes typified by low ecological connectivity and simplified arboreal structure limit resource availability and alter avian species composition and functional-trait distributions. This study analyzed the relationship between the taxonomic and functional diversity of bird assemblages across four tree-cover categories (open (OP), semi-open (SO), semi-closed (SC), and closed (CL)) using 491 point counts from 104 cattle-ranching landscape mosaics in the northwestern Colombian Amazon. A trait matrix was compiled for 342 recorded species distributed among four trophic guilds. Margalef, Shannon, and Simpson indices, together with five functional diversity indices, were calculated, and correlation coefficients between indices were estimated by guild and cover type. The tree-cover gradient differentially influenced guild-level richness, evenness, and functionality. In open and semi-closed covers, insectivores showed greater functional richness, with traits adapted to foraging in open habitats, whereas intermediate vegetation supported higher evenness and functional complementarity in frugivores and granivores. Closed covers imposed stronger filtering on trait diversity. The results are consistent with metacommunity dynamics in which local assemblages and functional spillover sustain ecological processes at the mosaic scale. For cattle-ranching landscape management, reducing habitat contrast within the productive matrix and strengthening forest connectivity are recommended to conserve species richness and ecosystem functions.

## Introduction

Tropical forests of the Amazon biome have undergone substantial transformations because of human settlement and the expansion of productive activities, particularly livestock production. In the Colombian Amazon, the area devoted to cattle-ranching systems increased from 14.6 to 24 million hectares over the last 50 years, accompanied by changes in land use and landscape configuration (Del Río Duque et al. 2022; Alvarado Sandino et al. 2023). This expansion leads to fragmentation of natural landscapes into patches of varied shapes and sizes, with a high degree of disturbance and increasing isolation within a homogeneous, high-contrast matrix dominated by pastures (Pérez-Cabral et al. 2021). According to Curtis et al. (2021) and de Souza Leite et al. (2022), human alteration of natural landscapes drives biodiversity loss while simplifying vegetation structure and altering faunal assemblages. By contrast, tree cover regulates microclimatic conditions, provides resources such as food and shelter, and enhances connectivity among patches (Bain et al. 2020; Anderle et al. 2023). The distribution of fauna in transformed landscapes is conditioned by the availability and spatial arrangement of resources (Davis et al. 2024), which are mediated by interspecific interactions that influence local diversity and community structure (Lowney and Thomson 2021).

Several studies have evaluated the influence of the expanding livestock frontier on the loss of biological diversity, particularly of avifauna (Alvarado Sandino et al. 2023; Giraldo et al. 2024; Díaz-Cháux et al. 2025a). Owing to their wide distribution, environmental sensitivity, and ecological functions, birds constitute a model biological group for assessing the effects of land-use change (Maas et al. 2015; Echeverri et al. 2020; Low et al. 2024). According to Peña et al. (2023), the magnitude of these responses varies as a function of species’ functional traits, which determine their capacity to adapt to local conditions across scales by promoting niche segregation and reducing interspecific competition (Bain et al. 2020; Molina-Marin et al. 2022). This functional variability underpins, among other outcomes, contributions to regulating ecosystem services such as seed dispersal, pollination, and biological pest control (Enríquez-Acevedo et al. 2020; Matuoka et al. 2020; Pérez-Cabral et al. 2021; Smith et al. 2022; Biswas et al. 2023; Figueroa-Alvarez et al. 2024).

In this context, bird assemblages make important contributions to sustaining ecological processes in productive landscapes (Velasquez-Valencia and Bonilla-Gomez 2019; Velásquez-Trujillo et al. 2021). Nevertheless, most studies on the effects of land-use change focus on the taxonomic dimension of diversity (Bełcik et al. 2020). In recent years, the functional approach has gained prominence because it enables assessment of how communities respond to environmental gradients and anthropogenic pressures (Biswas et al. 2023). Although a positive relationship between taxonomic and functional diversity has often been assumed (Xu et al. 2024), recent research indicates that such correspondence depends on habitat structural complexity and on species’ ecological traits, such as the degree of specialization (Boesing et al. 2021; Birch et al. 2024). According to Lee and Martin (2017) and de Souza Leite et al. (2022), within ranching mosaics denser covers act as environmental filters that constrain avian richness and abundance, whereas intermediate covers, characterized by greater heterogeneity and the presence of trees within the pasture matrix, promote species evenness and functional coexistence. From this perspective, bird assemblages in ranching landscapes conform to the metacommunity concept (Montoya 2021), whereby landscape-level environmental conditions determine the distribution of species diversity, leading to functional spillover at the mosaic scale that sustains key ecological processes (van Schalkwyk et al. 2020; Boesing et al. 2021; Montealegre-Talero et al. 2021; Béllo Carvalho et al. 2023). However, in the Colombian Amazon biome, knowledge remains limited regarding how this relationship varies across different types of tree cover within ranching landscapes and how it is expressed among specific ecological guilds.

This study aimed to analyze the relationship between the taxonomic and functional diversity of bird assemblages along a tree-cover gradient in cattle-ranching landscapes of the Colombian Amazon. It sought to answer the following research question: How does the tree-cover gradient influence the taxonomic and functional diversity of avian ecological guilds in these landscapes? The working hypothesis was that richness, evenness, and taxonomic diversity exhibit positive correlation patterns with functional diversity indices, modulated by cover type and by the trophic guild of the species.

The results show an overall, differentiated correlation pattern between taxonomic richness and evenness and the functional indices, particularly within the granivorous and insectivorous guilds. However, as habitat structural complexity increases—from open toward semi-closed or closed covers—these correlations tend to weaken or even reverse in more specialized guilds. This pattern reflects the action of environmental filtering processes that limit the presence of certain species, thereby promoting functional convergence and revealing potential ecological redundancies within bird assemblages. Taken together, these findings underscore that both habitat structure and the degree of trophic specialization significantly influence the relationship between the taxonomic and functional dimensions of biodiversity. Understanding these interactions is essential for identifying functional guilds that are sensitive to the loss of tree cover and for designing management and ecological restoration strategies that promote biodiversity resilience in tropical cattle-ranching landscapes.

## Materials and methods

### Study area

The study was carried out in cattle-ranching landscapes of the northwestern region of the Colombian Amazon, across twelve of the 16 municipalities of the department of Caquetá: Albania (1.22° N, 75.9119° W), Belén de los Andaquíes (1.434° N, 75.8477° W), El Doncello (1.662° N, 75.1033° W), El Paujl (1.4917° N, 75.1479° W), Florencia (1.531° N, 75.5647° W), La Montañita (1.3056° N, 75.2291° W), Milán (1.1719° N, 75.4231° W), Morelia (1.3921° N, 75.6695° W), Puerto Rico (1.8586° N, 75.0378° W), San José del Fragua (1.2712° N, 76.1255° W), San Vicente del Caguán (1.9163° N, 74.7259° W), and Solita (1.0985° N, 75.638° W), covering a total area of 1,336,931 ha (Fig. 1). The region exhibits a geomorphology dominated by rolling hills (lomerío), piedmont, and floodplains with slopes < 12% (Velasquez-Valencia and Bonilla-Gomez 2019). Total precipitation is 4,277 mm yr⁻¹ with a unimodal distribution and a peak rainy period from April to October. Mean temperature is 28.62 °C, and relative humidity is 86%.

**Fig. 1.**
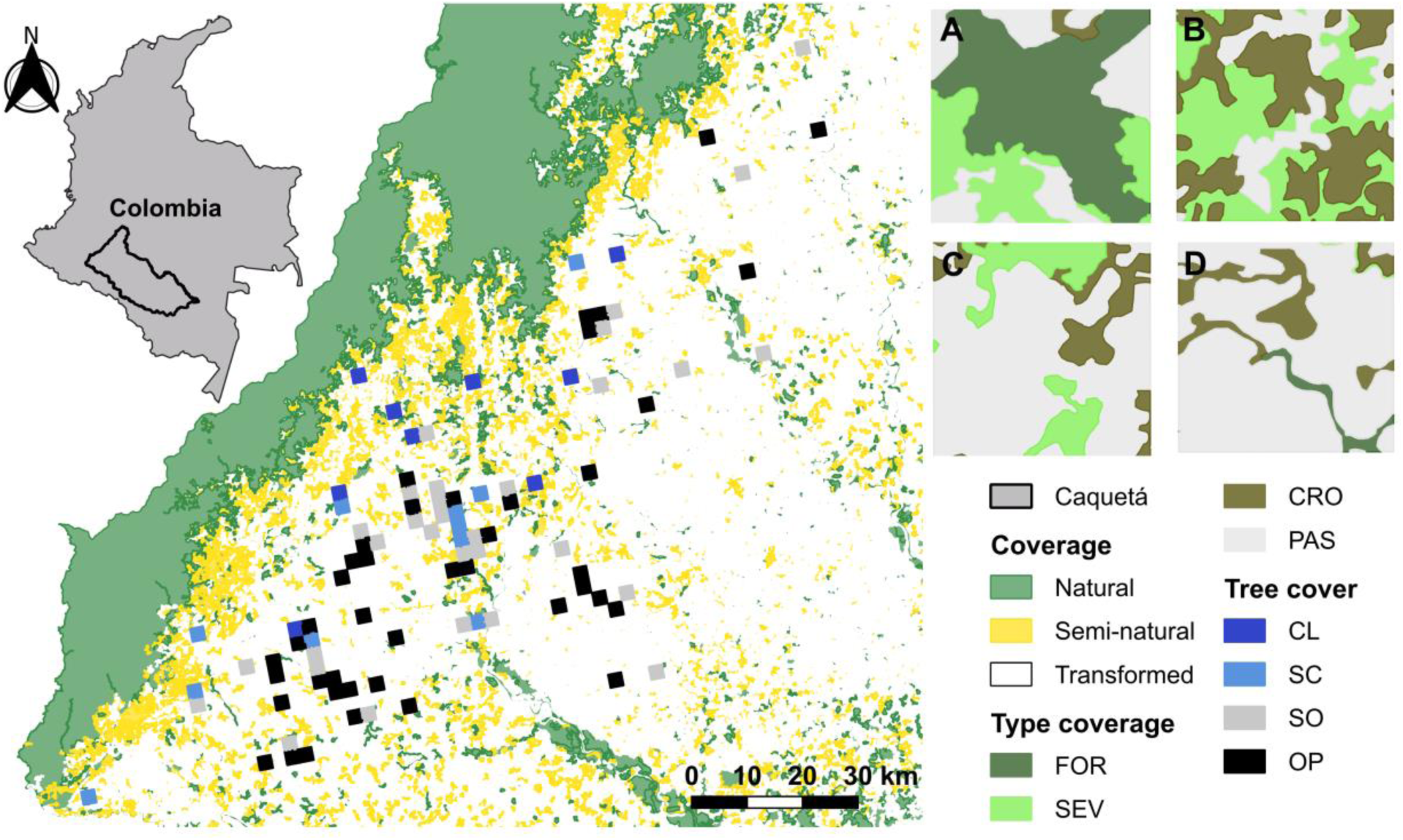
Study areas in the department of Caquetá with landscape units based on percentage of tree cover (black < 10%, gray 11–25%; light blue 26–45%; dark blue < 46%). (A) CL: closed; (B) SC: semiclosed; (C) SO: semiopen; (D) OP: open. FOR: forests; SEV: secondary vegetations; CRO: crops; PAS: pastures.

### Bird Census and Functional Traits

Bird diversity was surveyed between 2017 and 2023 using fixed-radius point counts conducted by a single observer (Alexander Velasquez-Valencia). Points were spaced at least 250 m apart, and all species seen or heard within a fixed 50 m radius at each point were recorded over 25 min. In total, 491 points were included, for a sampling effort of 12,276 observation minutes. Species-level taxonomy followed the South American Classification Committee (Remsen 2025), with verification against Colombian field guides (Echeverry-Galvis et al. 2022; Ayerbe Quiñones 2023; McMullan 2023). For fixed (life-history) traits, species were assigned to seven trophic guilds: frugivores (FRU), insectivores (INS), nectarivores (NEC), granivores (GRA), folivores (FOL), scavengers (CAR), and vertebrate consumers (VER) (Díaz-Cháux et al. 2025a).

The morphological trait matrix for identified species was built from measurements of specimens in the Ornithological Collection of the Natural History Museum of the Universidad de la Amazonia (UAM) and the global morphological, ecological, and geographic bird database AVONET (Tobias et al. 2022). The functional traits considered were total body length (LTO), tarsus length (LTA), wing chord (AEX), commissure (COM), bill height (ALT), culmen length (CTO), and body mass (PES).

### Classification of Mosaics in Cattle-Ranching Landscapes

Over the sampling extent (1,336,931 ha), a grid of 2.5 km × 2.5 km cells with a 0° rotation angle was generated in QGIS 3.40 (QGIS Development Team 2022) to align with the north–south axes under the national MAGNA-SIRGAS 2018 coordinate system (EPSG: 9377) and ensure spatial consistency. A total of 104 grid cells that contained point-count locations were selected for mosaic-scale analyses. Each selected mosaic covered 625 ha, a size that enhanced heterogeneity in the composition of vegetation covers present. According to Leyequién et al. (2010) and Martínez-Ramos et al. (2016), mosaics larger than 2 km² make it possible to reveal relationships between the diversity of functional guilds and qualitative landscape properties.

For each mosaic, a thematic land-cover map was produced based on the CORINE Land Cover methodology adapted for Colombia at 1:100,000 scale (IDEAM 2010) and for the Amazon region at 1:25,000 scale (Instituto SINCHI 2025). Spatial information was obtained from geospatial vector data downloaded from the geoportal of the Environmental Territorial Information System of the Colombian Amazon (SIAT-AC) for 2024, under the “Land Cover” category. Land-cover classification followed the procedure proposed by Velasquez-Valencia and Bonilla-Gomez (2019). This classification was cross-checked against Amazon-specific reference coverages (Díaz-Cháux et al. 2025b) to ensure accuracy and currency, and it was validated through direct field observations.

The mapped coverage patches were digitized and assigned to four categories: forests, secondary vegetation, croplands, and pastures. These data were converted to Shapefile format and georeferenced under the national MAGNA-SIRGAS 2018 coordinate system (EPSG:9377) using ArcMap 10.8 (Esri 2024). The Shapefile was then converted to a raster with a spatial resolution of 2 m per pixel in QGIS 3.40 (QGIS Development Team 2022). Landscape structure and composition were quantified for each mosaic with FragStat version 4.2 (McGarigal et al. 2024). Landscape metrics were selected for their ecological relevance, as they reflect habitat availability and connectivity—key aspects shaping avian community responses (Adler and Jedicke 2022; Ramos-Mosquera et al. 2025).

Mosaics were classified according to the area occupied by land-cover patches. Specifically, the total percentage of tree cover in each mosaic was calculated following IDEAM (2010) criteria, yielding four types that define the tree-cover gradient in cattle-ranching landscapes of the Colombian Amazon: (i) open mosaics (OP), with < 10% tree cover across the mosaic (grasslands/pastures); (ii) semi-open mosaics (SO), with < 25% tree cover (agroforestry systems); (iii) semi-closed mosaics (SC), with < 45% tree cover (secondary and riparian vegetation); and (iv) closed mosaics (CL), with >45% tree cover across the mosaic (Table S1 – Supplementary material).

### Statistical Analysis

A generalized linear model (GLM) analysis of variance was used with a Poisson error distribution to test for differences in the mean distribution of species richness and individual abundance of birds among the four tree-cover types that constitute the gradient in cattle-ranching landscapes of the Colombian Amazon. Overdispersion was evaluated via the ratio of model deviance to error degrees of freedom; when overdispersion was detected, the Poisson distribution was replaced with a negative binomial distribution. Pairwise mean comparisons were conducted with Fisher’s LSD test (α = 0.05). Analyses were performed in Navure version 3.1.0 (Navure Team 2025).

To assess sampling integrity/behavior and the completeness of species richness across mosaics, species accumulation curves were plotted using nonparametric richness estimators Chao1, Chao2, first-order Jackknife, and second-order Jackknife (Gotelli and Chao 2013; Chao and Chiu 2016), based on the records from the 104 tree-cover mosaics. These were computed with rarefaction and accumulation functions in the vegan 2.8-0 package (Oksanen et al. 2025) for R version 4.2 (R Core Team 2022). Community abundance structure was evaluated by fitting species-abundance distribution models to the data pooled across all cover types and separately by cover (OP, SO, SC, CL). Goodness of fit of empirical data to theoretical models was tested using chi-square tests, implemented in PAST (PAleontological STatistics) version 4.16 (Hammer et al. 2001; Hammer 2024).

Alpha diversity was estimated using Hill numbers, expressing the effective number of species for the total sample and by guild within each tree-cover type. Rarefaction–extrapolation curves were plotted for the first three Hill numbers: q0 (species richness), q1 (Shannon diversity), and q2 (Simpson dominance) (Chao et al. 2014), using the iNEXT.3D package (Chao et al. 2021) in R version 4.2 (R Core Team 2022).

Functional diversity of the bird community across tree-cover types was quantified with five multidimensional indices (Pla et al. 2012; Bonfim et al. 2021): functional richness (FRic), functional evenness (FEve), functional divergence (FDiv), functional dispersion (FDis), and Rao’s quadratic entropy (RaoQ), using the FD package (Laliberté et al. 2014) in R version 4.2 (R Core Team 2022). An analysis of variance with linear models (LM) was used to test for differences in taxonomic diversity indices and functional diversity indices among cover types and among guilds (GRA, INS, FRU, VER) within each cover type. Response variables were the three taxonomic indices (Simpson, Margalef, Shannon) and the five functional indices (FRic, FEve, FDiv, FDis, RaoQ); fixed effects were the four tree-cover types (OP, SO, SC, CL). The same analysis was performed separately for each of the four guilds. Fisher’s LSD test (α = 0.05) was used for mean comparisons, and analyses were conducted in InfoStat version 2020 (Di Rienzo et al. 2020).

To evaluate the relationship between taxonomic and functional diversity in bird communities, Pearson correlation coefficients were calculated by guild and tree-cover type (Flynn et al. 2009). Assumptions of normality and linearity were verified beforehand with Shapiro–Wilk tests and scatterplots. Additionally, a principal components analysis (PCA) was performed using taxonomic and functional indices as variables to graphically depict association patterns among guilds and tree-cover types via a biplot. All analyses were conducted in InfoStat version 2020 (Di Rienzo et al. 2020).

## Results

A total of 16,244 individual birds were recorded, distributed across 342 species, 52 families, and 25 orders (Table S2 — Supplementary material). Of all individuals sampled, 38.63% were concentrated in ten species belonging to eight families. Tyrannidae showed the greatest richness and abundance, with 65 species and 2,312 individuals. Mean richness and abundance did not differ significantly among cover types (p > 0.05). Nonetheless, a declining trend in the number of species and individuals was observed along the tree-cover gradient. Open covers yielded the highest values (6,750 individuals, 248 species, 47 families, 21 orders), followed by semi-closed (4,236 individuals, 227 species, 47 families, 23 orders) and semi-open covers (2,882 individuals, 190 species, 45 families, 19 orders). Closed areas had the lowest abundance and richness (2,376 individuals, 188 species, 45 families, 20 orders).

Abundance distributions conformed to the geometric model in the overall sample (k = 0.0172; χ² = 1.34 × 10^^−4^; p < 0.0001) and within semi-open (k = 0.0228; χ² = 996.2; p < 0.0001), semi-closed (k = 0.0193; χ² = 1,827; p < 0.0001), and closed covers (k = 0.0230; χ² = 2,443; p < 0.0001), indicating few dominant species and a higher proportion of low-abundance species. In contrast, the open cover fitted a log-normal distribution (log-normal mean = 0.7957; variance = 0.5732; χ² = 17.84; p = 0.0127), in which most species had intermediate abundance, and few were highly abundant. The most numerous and widespread species across mosaics and cover types were *Ardea ibis* (1,145 individuals), *Ara severus* (922), *Crotophaga ani* (874), and *Thraupis episcopus* (620).

Observed species richness in the 104 evaluated mosaics represented more than 88% of the richness estimated by nonparametric estimators (Fig. 2). Accordingly, the sampling effort—as reflected in recorded bird richness— provides a representative picture of regional diversity; the analyses therefore capture diversity patterns and help elucidate the ecological processes structuring bird communities in Amazonian cattle-ranching mosaics.

**Fig. 2.**
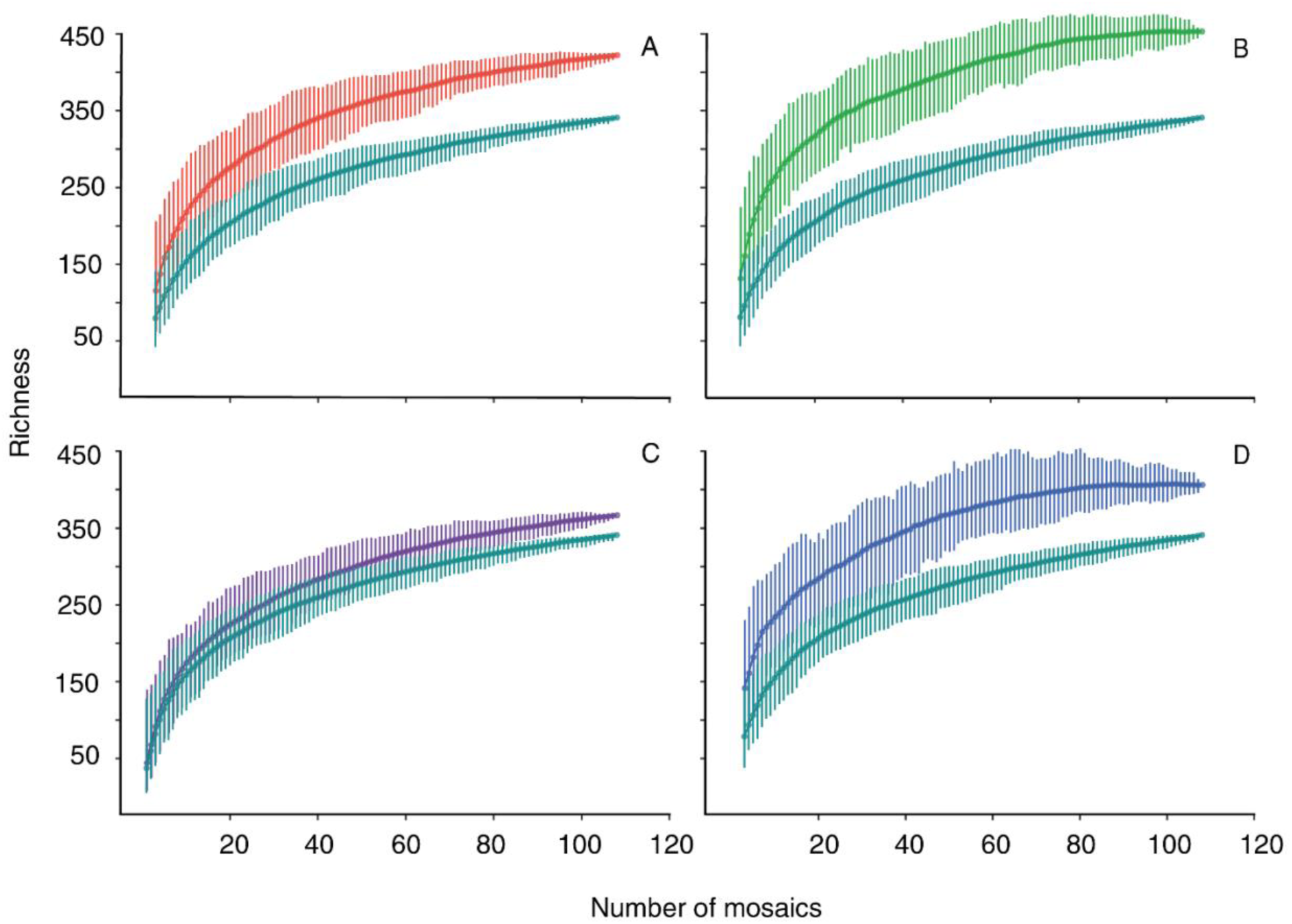
Species accumulation curves for birds across tree-cover types in cattle-ranching landscapes of the Colombian Amazon. Each sampling unit comprises 104 landscape mosaics. Chao 1 (A), Chao 2 (B), first-order Jackknife (C), second-order Jackknife (D).

For Hill numbers, at q0 (richness) the open cover (OP) showed the greatest number of species, followed by the semi-closed (SC) cover. Likewise, OP exhibited the highest Shannon (q1) and Simpson (q2) diversity relative to the other covers (Fig. 3A). Sample completeness is interpreted as the number of individuals present at a site divided by the number of individuals observed at that site. In this regard, all covers reached completeness; however, semi-open (SO) and closed (CL) covers attained completeness with the fewest individuals, whereas OP reached completeness only after accumulating roughly three times the number of individuals recorded for SO and CL (Fig. 3B).

**Fig. 3.**
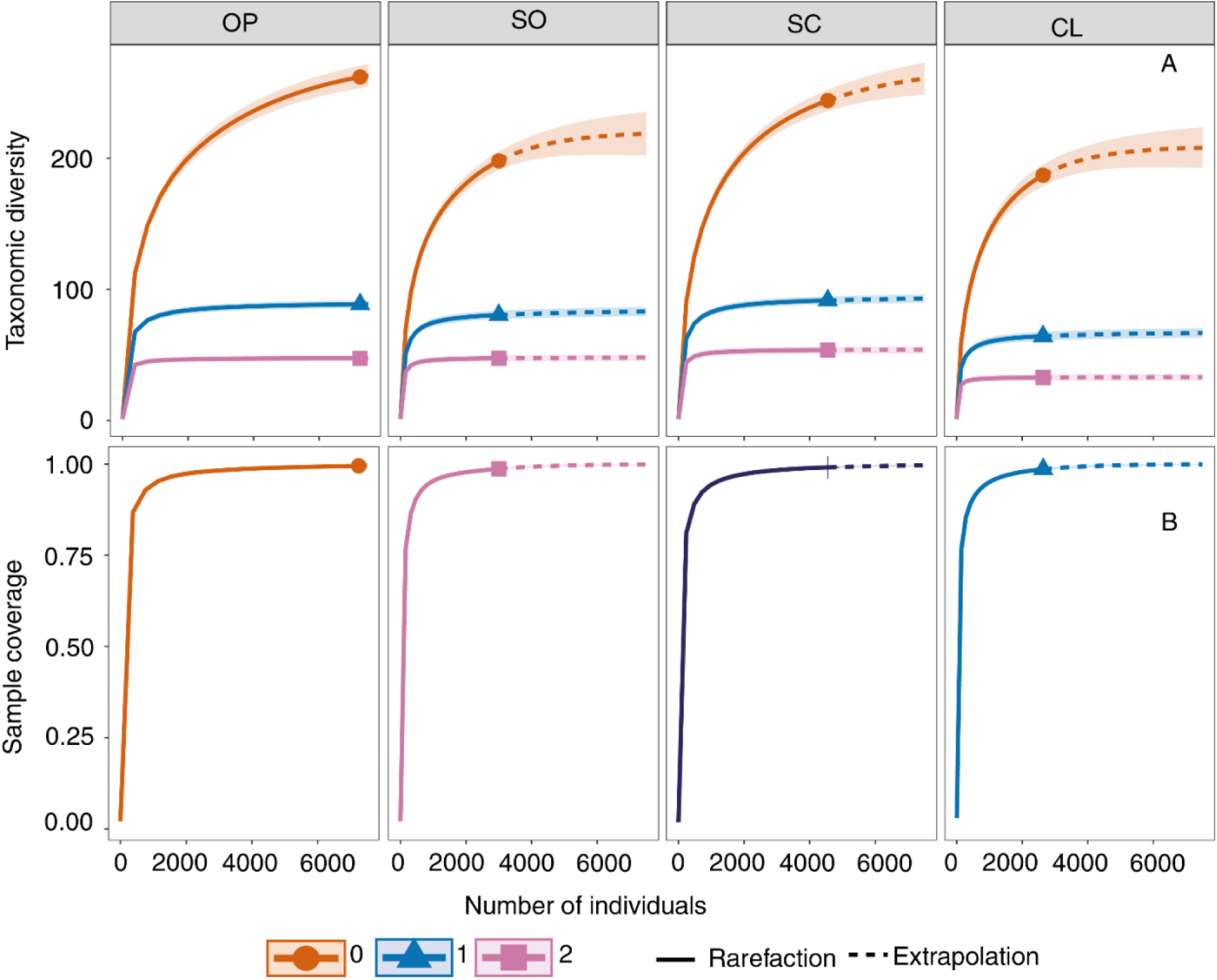
Rarefaction–extrapolation curves for diversity of order q0, q1, and q2 for bird species across tree-cover types in cattle-ranching landscapes of the Colombian Amazon. SO: semi-open; OP: open; SC: semi-closed; CL: closed. (A) Sample-size–based sampling curve; (B) Sample completeness curve.

### Taxonomic and functional diversity of the bird assemblage by cover types

Mean values of taxonomic and functional diversity indices across the different tree-cover types evaluated in cattle-ranching landscapes were not statistically different (p > 0.05). Overall, a positive and significant correlation was found between the taxonomic diversity indices of Shannon and Margalef and the functional diversity indices FDis, FRic, and RaoQ (Table 1). By contrast, the Simpson index correlated negatively with FDis, FRic, RaoQ, and FEve. Functional divergence (FDiv) showed no associations with taxonomic diversity.

**Table 1.**
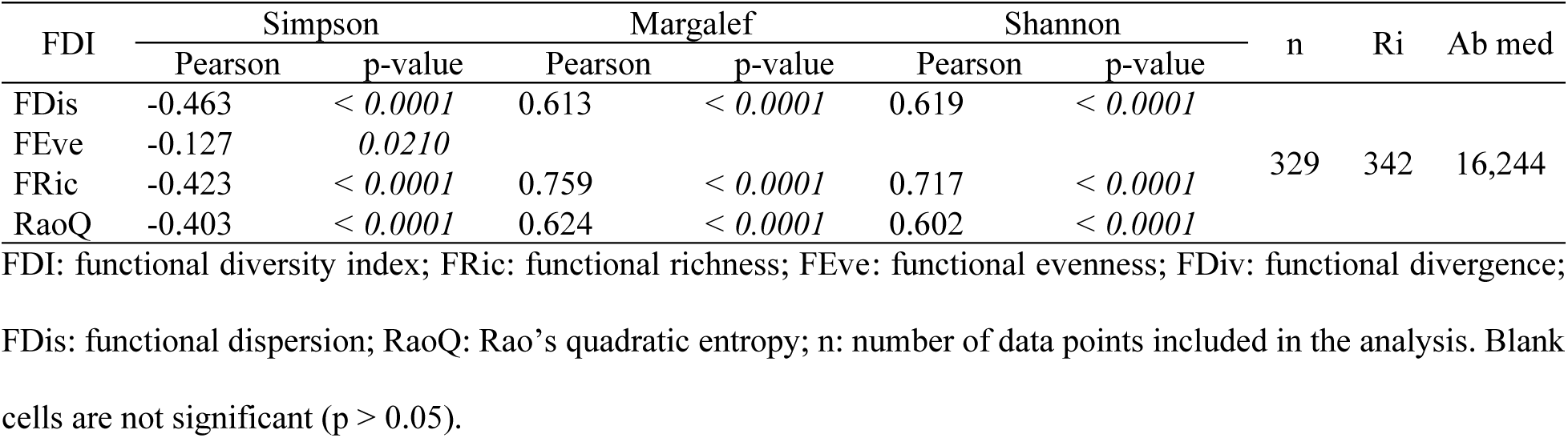
Pearson correlation coefficients between taxonomic diversity indices and functional diversity indices for the bird assemblage across tree-cover types in cattle-ranching landscapes of the Colombian Amazon.

| FDI | Simpson |  | Margalef |  | Shannon |  | n | Ri | Ab med |
| --- | --- | --- | --- | --- | --- | --- | --- | --- | --- |
|  | Pearson | p-value | Pearson | p-value | Pearson | p-value |  |  |  |
| FDIs | -0.463 | < 0.0001 | 0.613 | < 0.0001 | 0.619 | < 0.0001 | 329 | 342 | 16,244 |
| FEve | -0.127 | 0.0210 |  |  |  |  |  |  |  |
| FRic | -0.423 | < 0.0001 | 0.759 | < 0.0001 | 0.717 | < 0.0001 |  |  |  |
| RaoQ | -0.403 | < 0.0001 | 0.624 | < 0.0001 | 0.602 | < 0.0001 |  |  |  |

FDI: functional diversity index; FRic: functional richness; FEve: functional evenness; FDiv: functional divergence; FDis: functional dispersion; RaoQ: Rao's quadratic entropy; n: number of data points included in the analysis. Blank cells are not significant ( $p > 0.05$ ).

A consistent pattern across vegetation covers was observed in the correlations between functional and taxonomic diversity indices: the functional indices FDis, FRic, and RaoQ showed significant positive correlations with Shannon and Margalef, and negative correlations with Simpson (Table 2). Nevertheless, in open and closed covers, functional diversity exhibited a stronger correlation with Margalef species richness, whereas in semi-open and semi-closed covers the strongest correlation was with the Shannon diversity index.

**Table 2.**
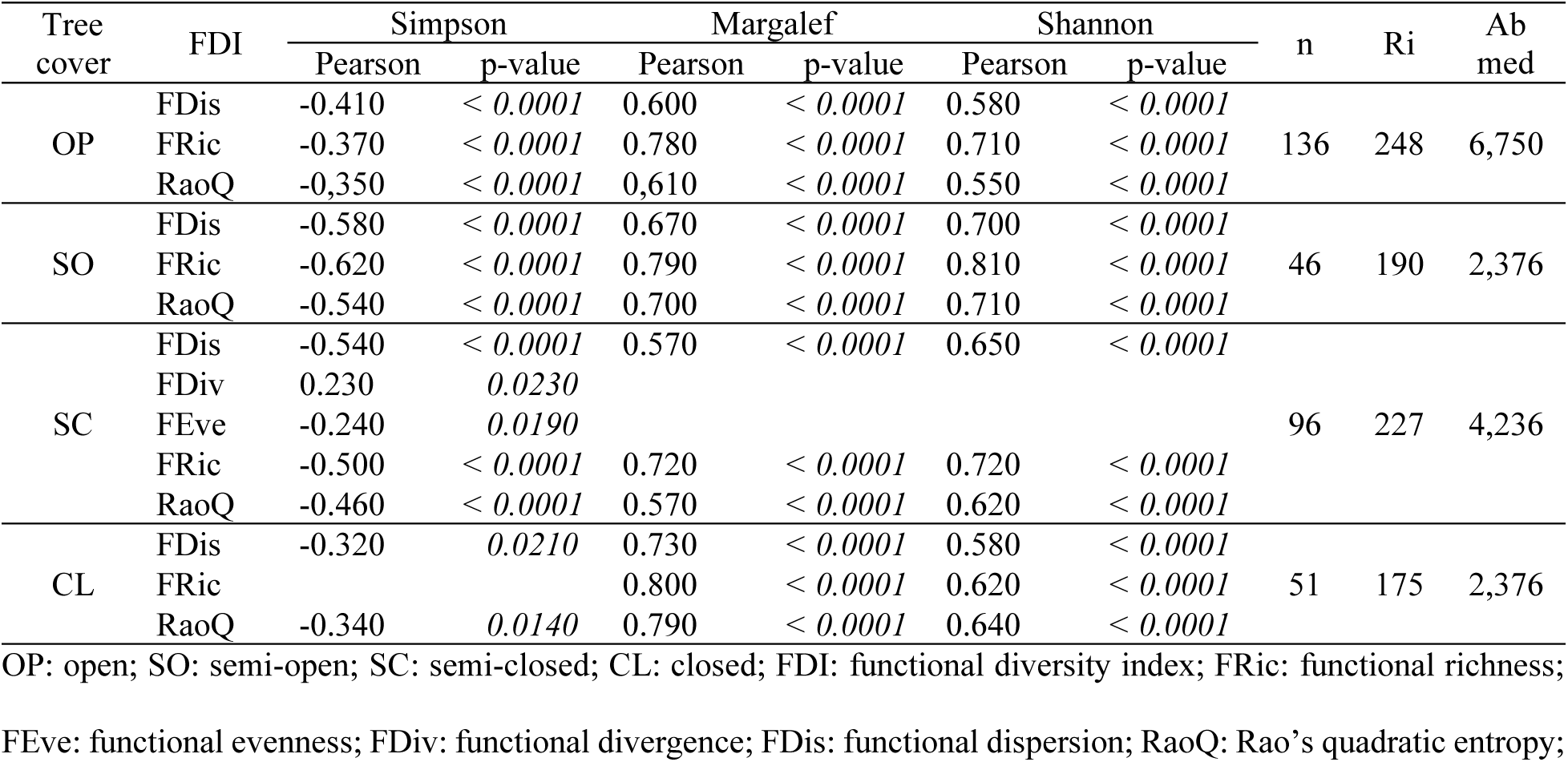
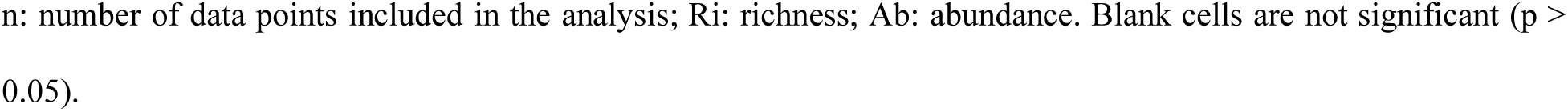
Pearson correlation coefficients between taxonomic and functional diversity indices of the bird assemblage, compared among tree-cover types in cattle-ranching landscapes of the Colombian Amazon.

### Taxonomic and functional diversity by guilds across cover types

No significant differences were detected in mean richness and abundance within guilds across cover types, but there were differences among guilds within each tree-cover type (p < 0.05). Insectivores (INS) exhibited the highest richness and abundance in all cover types, with peak values in open cover (Table S3, Supplementary material). In the remaining cover types, INS abundance did not differ from that of frugivores (FRU) but did differ from vertebrate consumers (VER) and granivores (GRA). Closed cover showed the lowest taxonomic diversity values for all guilds (Fig. 4). Richness (q0) was highest for FRU and GRA in semi-closed cover, whereas for INS and VER it was highest in open cover. All guilds displayed higher Shannon (q1) and Simpson (q2) diversity in semi-closed cover.

**Fig. 4.**
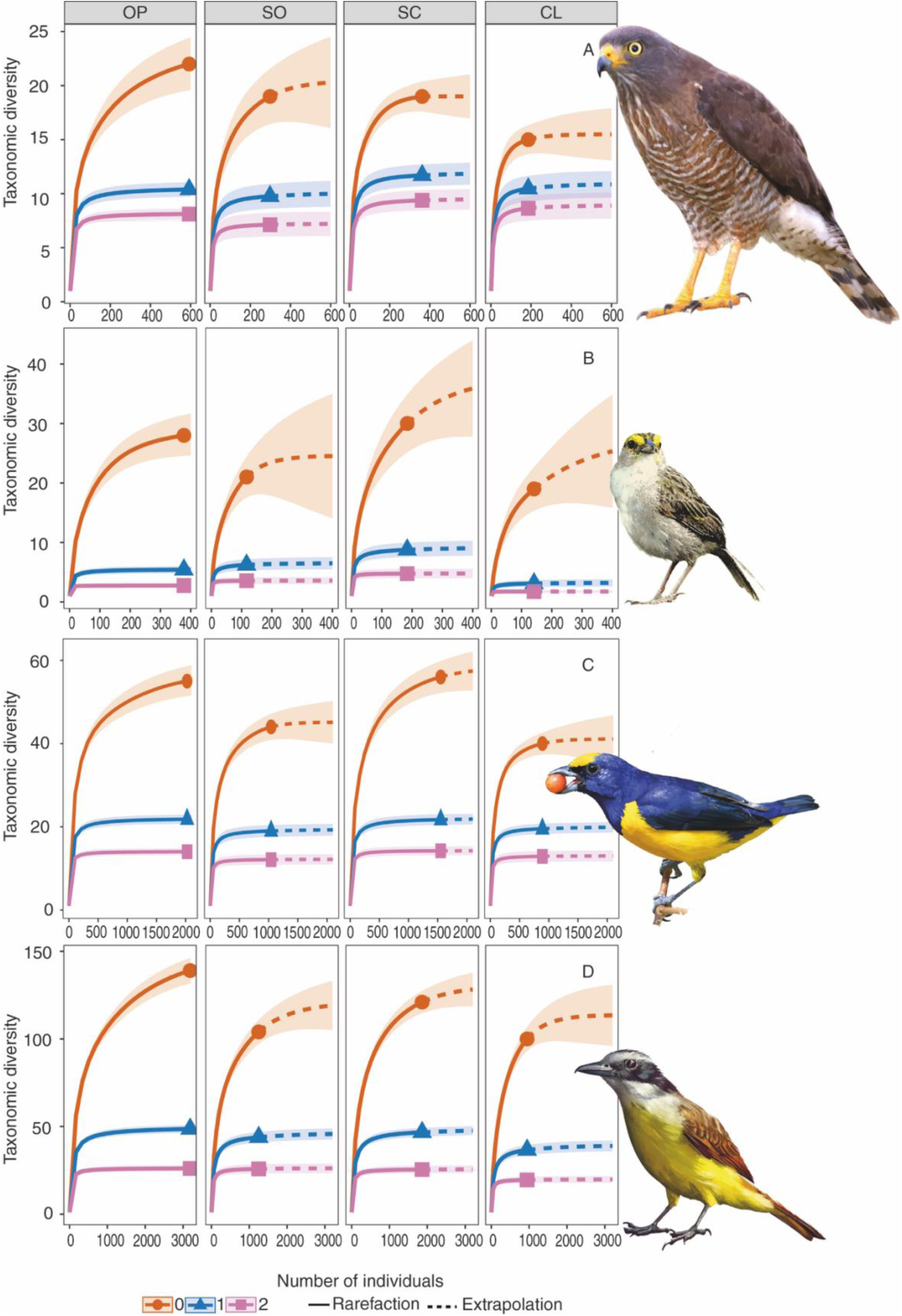
Rarefaction–extrapolation curves of q0, q1, and q2 diversity for bird guilds across tree-cover types in cattle-ranching landscapes of the Colombian Amazon. SO: semi-open; OP: open; SC: semi-closed; CL: closed. (A) VER: vertebrate consumers; (B) GRA: granivores; (C) FRU: frugivores; (D) INS: insectivores.

Functional diversity indices differed significantly (p < 0.05) among guilds within each tree-cover type (Supplementary material S3). Insectivores (INS) showed the highest FDis values in all covers, especially in semi-open cover. Functional divergence (FDiv) was greater for FRU and GRA in semi-closed cover, and for INS in open and semi-open covers. For functional evenness (FEve), no differences among guilds were detected in semi-open cover; in the other covers, INS, GRA, and VER did not differ from one another but did differ from FRU. For functional richness (FRic), INS and FRU had the highest values within covers, and these among-guild differences were significant. Rao’s quadratic entropy (RaoQ) was highest for INS and lowest for VER in all cover types.

In the principal component analysis (PCA) of taxonomic and functional diversity indices by guild and cover, the first two components explained 50% of the total variability (Fig. 5). Along PC1, insectivores grouped on the positive side with the taxonomic indices Shannon and Margalef and with the functional indices FDis, FRic, and RaoQ. On the same axis, frugivores were positively associated with FDiv and FEve, chiefly in semi-closed cover. Granivores and vertebrate consumers were positioned on the negative side and associated with the Simpson index. Along PC2, INS, FRU, and VER grouped with RaoQ, FDis, FDiv, FEve, and Simpson in covers of intermediate to higher structural complexity (SO, SC, and CL), whereas GRA occupied the negative end, associated with FRic, Shannon, and Margalef in open cover.

**Fig. 5.**
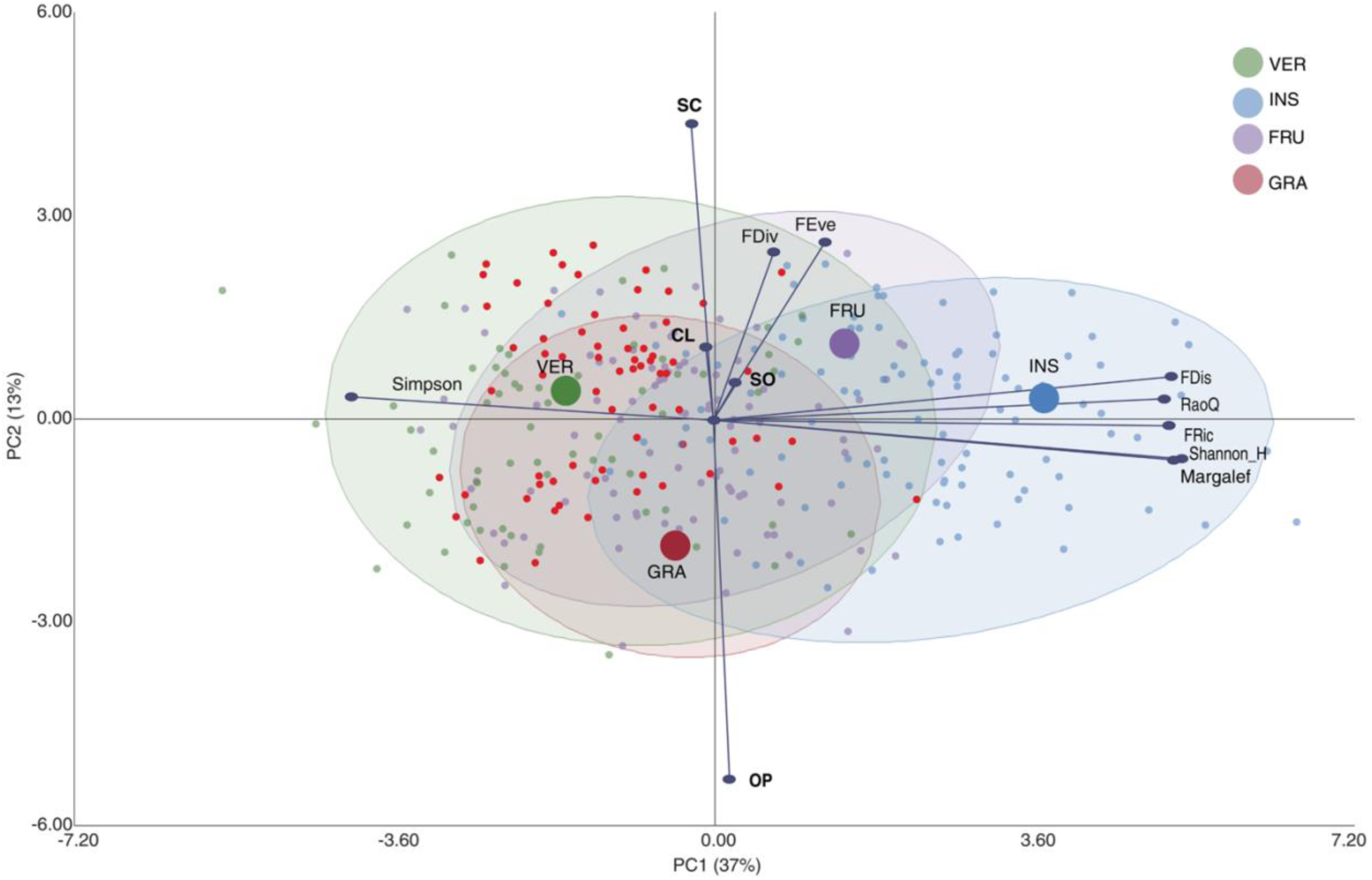
Biplot from principal component analysis (PCA) of taxonomic and functional diversity indices for guilds across tree-cover types in cattle-ranching landscapes of the Colombian Amazon. SO: semi-open; OP: open; SC: semi-closed; CL: closed; INS: insectivores; FRU: frugivores; GRA: granivores; VER: vertebrate consumers; FRic: functional richness; FEve: functional evenness; FDiv: functional divergence; FDis: functional dispersion; RaoQ: Rao’s quadratic entropy.

The relationship between taxonomic and functional diversity in cattle-ranching landscapes varied with tree-cover typology and ecological guild. A general pattern of positive correlation was identified between taxonomic diversity (Margalef and Shannon) and functional diversity (FRic, FDis, RaoQ, FEve) in bird communities (Table 3). By contrast, Simpson dominance and the functional indices—except FDiv—showed negative correlations across guilds and cover types.

**Table 3.**
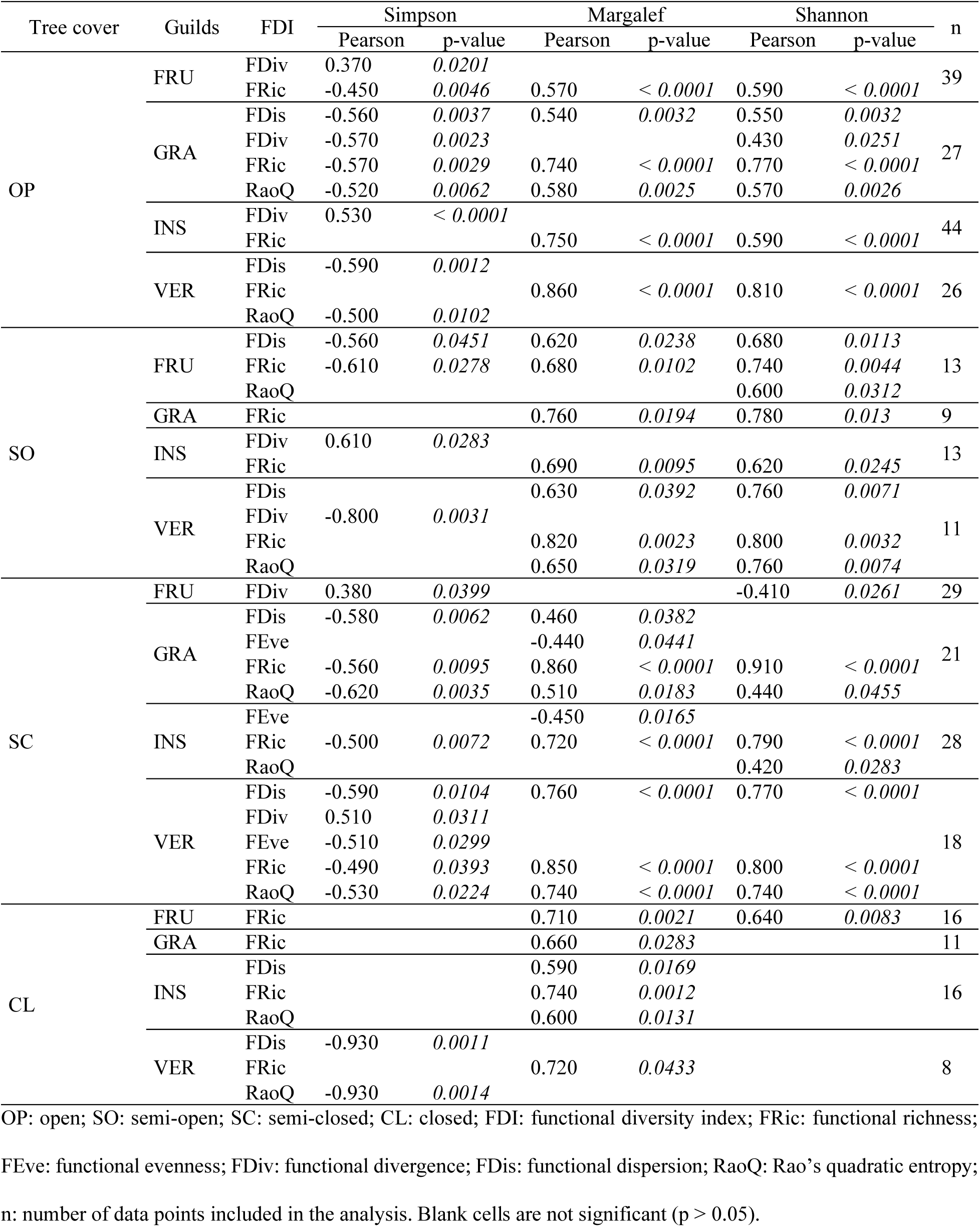
Pearson correlation coefficients between taxonomic and functional diversity indices by guild across tree-cover types in cattle-ranching landscapes of the Colombian Amazon. All covers with all guilds, each cover with all guilds, and each cover with each guild were analyzed.

In open and semi-open covers, positive correlations predominated for frugivores, insectivores, and granivores; granivores in particular showed stronger positive correlations between Shannon/Margalef and all functional indices, and negative correlations with Simpson. In open covers, insectivores and frugivores displayed positive correlations between Simpson and FDiv, but in semi-open covers Simpson correlated negatively with FRic and FDis.

In closed covers, positive correlations were observed between Margalef and FRic across all guilds, whereas Shannon correlated with FRic only for frugivores. The insectivore guild showed positive correlations between FDiv– RaoQ and the taxonomic indices, while vertebrate consumers exhibited negative correlations between Simpson and FDis–RaoQ. Overall, FRic and RaoQ were the most sensitive to variation in taxonomic richness, whereas FDiv was differentially associated with dominance.

## Discussion

The bird species recorded in this study correspond exclusively to records from the cattle-ranching landscapes of the Colombian Amazon and represent roughly 20% of the avifauna reported for Colombia (1,821 spp.) and 33.27% of that reported for the department of Caquetá, in the northwestern Amazon region (1,028 spp.) (Rheindt et al. 2025; SiB Colombia 2025). The observed richness indicates high sampling completeness and efficient species detection for the study area. These results support the robustness of the inventory and provide a reliable basis for analyzing ecological patterns such as the diversity and functionality of bird communities in livestock-production mosaics.

For the Colombian Amazon, this study constitutes the first effort to demonstrate how landscape structure— evaluated through a gradient of tree cover—shapes the relationship between the taxonomic and functional dimensions of the avifauna. In line with studies from the region (Díaz-Cháux et al. 2025a; Díaz-Cháux et al. 2025b), understanding these interactions helps identify functional guilds that are sensitive to habitat disturbance and informs the design of management and ecological restoration strategies to strengthen biodiversity resilience in productive tropical landscapes.

### Taxonomic and functional diversity of the bird assemblage across cover types

The taxonomic diversity of the bird assemblage responded divergently to changes in tree density; the ranching matrix acts as an environmental filter. This condition favors communities composed of generalist bird species with low habitat requirements, benefiting from low spatial heterogeneity (Boesing et al. 2021; Birch et al. 2024; Xu et al. 2024). In this regard, closed covers exhibited low richness and evenness, with dominance by a few species of low ecological specialization and a higher proportion of rare species dependent on specific microhabitats (de Souza Leite et al. 2022). By contrast, open areas supported more diverse and even communities, with the coexistence of species of intermediate abundances adapted to disturbed environments (Alba et al. 2025), such as *Ardea ibis*, *Ara severus*, *Crotophaga ani*, and *Thraupis episcopus*, whose presence is explained as the result of spillover from nearby, more heterogeneous patches (Montealegre-Talero et al. 2021; de Souza Leite et al. 2022; Béllo Carvalho et al. 2023; Carvalho et al. 2023). These results are consistent with other reports from Amazonian landscapes, where structural simplification favors species with broad ecological tolerance and geographic distributions (Neate-Clegg and Şekercioǧlu 2020; Eyster et al. 2022).

According to Siqueira et al. (2025), species richness and the evenness of abundances determine the variety of functions and the breadth of the ecological niche. More diverse communities occupy a wider functional space and exhibit greater ecological stability (Dardanelli et al. 2022). As dominance increases, functions become concentrated in a few species and functional redundancy declines (Low et al. 2024), thereby increasing ecosystem vulnerability (Luck et al. 2013). In the ranching landscapes studied here, the relationship between taxonomic and functional diversity was modulated by the tree-cover gradient (Rurangwa et al. 2022; Díaz-Cháux et al. 2025b). At the extremes of the gradient, despite differences in the taxonomic dimension, open and closed covers exhibited a strong positive correlation between species richness and functional dispersion. As noted by Wei et al. (2025), in these areas the addition of new species expands functional space, reduces redundancy, and increases niche complementarity. This pattern reflects processes mediated by environmental filters (van Schalkwyk et al. 2020) and metacommunity dynamics (Montoya 2021), whereby differential colonization and the spillover of species and individuals from adjacent habitats sustain functionally diverse assemblages. Intermediate covers simultaneously maximize richness and evenness, with a more balanced functional distribution (Gantz et al. 2024; Tscharntke et al. 2025), in which functionally similar species enhance stability and promote the retention of communities’ functional diversity (Echeverri et al. 2020; Giraldo et al. 2024; Kiat and O’Connor 2024).

### Taxonomic and Functional Diversity by Guilds Across Cover Types

Taxonomic diversity varied among guilds along the tree-cover gradient; insectivores exhibited higher richness and abundance in open areas. This guild benefits from the high availability of arthropods at edges and in well-lit understories, which favors species with active foraging strategies (Romero-Díaz et al. 2020; Jarrett et al. 2021; Matyjasiak et al. 2023; Ayomiposi Ayodele 2025). Semi-closed cover types showed greater evenness among guilds and high richness of frugivores and granivores, indicating that intermediate heterogeneity promotes functional coexistence. By contrast, closed cover types recorded the lowest values, likely associated with the high structural specialization of communities constrained by limited resource availability (Joyce et al. 2024; Restrepo-Cardona et al. 2024).

The tree-cover gradient differentially influences the taxonomic and functional diversity of bird guilds in ranching landscapes, yielding a pattern consistent with environmental filtering processes (Codesido and Bilenca 2021; Niklison et al. 2024; Alba et al. 2025). Insectivores showed the highest functional dispersion, especially in semi-open cover types, indicating broader niche breadth and ecological plasticity. These are traits that favor this guild’s association with fragmented areas and low forest dependence (Romero-Díaz et al. 2020; Ugalde-Lezama et al. 2022; Figueroa-Alvarez et al. 2024).

Frugivores, for their part, maintained high richness and functional evenness across the gradient, contributing to processes such as seed dispersal and vegetation regeneration that are essential for the recovery of degraded areas (Díaz Vélez et al. 2015; Maya-Girón et al. 2023; Hasui et al. 2024). Greater functional divergence of frugivores and granivores in semi-closed cover types, and of insectivores in open areas, indicates that structural heterogeneity not only promotes coexistence but also secures functional complementarity (Levey et al. 2025).

In open and semi-closed cover types, the higher richness and functional dispersion of insectivores reflects selection for species with traits adapted to foraging in low-density habitats (Silva et al. 2020; Gaston 2022; Téllez-Hernández et al. 2023; Guerrero et al. 2024), whereas in intermediate cover types frugivores and granivores exhibit greater evenness and functional divergence, evidencing complementary functional roles associated with structural heterogeneity (Posso et al. 2024). By contrast, closed cover types act as more restrictive filters by limiting diversity in both dimensions (Lee and Martin 2017; Nogueira et al. 2021; Hua et al. 2024; Fontana et al. 2025). These results suggest that bird communities are organized as metacommunities in which local assemblages respond to differential resource availability and habitat structure, while simultaneously generating functional spillover at the mosaic scale and ensuring the persistence of key ecological processes within livestock matrices of the Colombian Amazon.

### Implications for Management and Ecological Restoration

The findings of this study have direct implications for the management of ranching landscapes in the Colombian Amazon, as they show that the taxonomic and functional diversity of birds responds differentially to the tree-cover gradient. In line with Ocampo-Ariza et al. (2024), these patterns indicate that bird assemblages are structured as metacommunities, whose dynamics reveal the multidimensionality and spatial dependence of diversity.

The high representativeness of the recorded avifauna indicates that, despite transformation, these systems retain substantial conservation potential if appropriately managed. Structural heterogeneity emerges as a key factor: semi-closed cover types sustain the greatest functional complementarity among guilds, whereas open and closed areas contribute differently to the provision of ecosystem functions (Farneda et al. 2025). This pattern suggests that management strategies should prioritize heterogeneous mosaics with high structural and functional connectivity, integrating patches under natural regeneration, biological corridors, and silvopastoral systems, with the objective of maintaining both species richness and functional resilience, as well as the provision of important regulating ecosystem services (Alvarado Sandino et al. 2023). According to Oropeza-Sánchez et al. (2025), higher forest richness increases bird–plant interactions, which promotes fruit consumption and, consequently, the regeneration of fragmented areas (Pizo et al. 2021). Accordingly, identifying guilds—such as frugivores dependent on intermediate cover—underscores the importance of maintaining dispersal and vegetation succession processes (Béllo Carvalho et al. 2023; Carvalho et al. 2023). In this sense, ranching landscapes should not be regarded solely as areas of biodiversity loss, but as strategic settings where appropriate management can mitigate impacts, strengthen ecological functions, and contribute to regional sustainability.

## Conclusions

In ranching landscapes of the Colombian Amazon, the tree-cover gradient modulates the relationship between the taxonomic and functional diversity of bird communities, underscoring the importance of integrating both dimensions to understand assemblage dynamics in productive systems. The positive correlation between richness and functional dispersion at the extremes of the gradient indicates that species additions expand functional space and promote niche complementarity, a pattern consistent with environmental filtering mechanisms and spillover dynamics from adjacent habitats. These findings reinforce the value of functional diversity as a metric for assessing ecological resilience, as it captures the responses of guilds to landscape structural heterogeneity and habitat disturbance with precision. In this regard, reducing contrast within the productive matrix and promoting connectivity among forest fragments emerge as priority strategies to maintain essential ecosystem functions and reduce extinction risk for specialist species and guilds. Consequently, the integration of taxonomic and functional indices should be incorporated as a strategic benchmark in monitoring, restoration, and the planning of conservation units in Amazonian ranching landscapes.

## Statements and declarations

### Author contributions

AVV realizó el censo de aves, JTD, AVV y ANV realizaron la conceptualización, investigación, metodología, curación de datos y análisis formal. LPG gestionó parte de la financiación del estudio. ANM elaboró la clasificación de los mosaicos y el mapa temático de coberturas. AVV y JTD, construyeron el primer borrador del manuscrito y MAB y FC realizaron la revisión y evaluación crítica del contenido intelectual del documento. Todas las versiones del manuscrito fueron revisadas y validadas por todos los autores. Todos los autores leyeron y aprobaron el manuscrito final.

### Ethics statement

The specimen collections were conducted under the framework of Resolution 1015 of 2016, in accordance with the General Permit for the Collection of Specimens of Wildlife Species for Non-Commercial Scientific Research, issued to the Universidad de la Amazonia. Approval for practices involving the use of animals was granted by the Institutional Committee on Ethics and Bioethics in Research of the Universidad de la Amazonia (FO-A-APC-01-11), established through Agreement 020 of 2018 by the Superior Council.

