## Supplementary material for "Tree-Cover Gradients Modulate the Taxonomic and Functional Diversity of Birds in Amazonian Cattle-Ranching Landscapes": Table S1

**Table S1.** Percentage gradient of tree cover in Amazonian livestock landscape mosaics.

| **id** | **Mosaics** | **Tree cover percentage gradient** | | | | **Latitude** | **Longitude** |
| --- | --- | --- | --- | --- | --- | --- | --- |
|  |  | **OP** | **SO** | **SC** | **CL** |  |  |
| 1 | M_001 |  |  |  | 46.4% | 1.146635 | - 76.198618 |
| 2 | M_003 |  |  |  | 47.4% | 1.272402 | - 76.006476 |
| 3 | M_004 |  | 13.0% |  |  | 1.265434 | - 76.006326 |
| 4 | M_005 |  |  |  | 44.6% | 1.370068 | - 75.984050 |
| 5 | M_006 |  | 12.2% |  |  | 1.302549 | - 75.909386 |
| 6 | M_007 | 8.3% |  |  |  | 1.146436 | - 75.914199 |
| 7 | M_009 | 6.6% |  |  |  | 1.293370 | - 75.871697 |
| 8 | M_010 |  | 10.7% |  |  | 1.273029 | - 75.873920 |
| 9 | M_011 | 7.4% |  |  |  | 1.227837 | - 75.869380 |
| 10 | M_012 |  | 10.6% |  |  | 1.155523 | - 75.867060 |
| 11 | M_013 | 9.1% |  |  |  | 1.138484 | - 75.872837 |
| 12 | M_014 |  | 14.8% |  |  | 1.139638 | - 75.853813 |
| 13 | M_015 |  |  |  | 45.0% | 1.338326 | - 75.822768 |
| 14 | M_016 | 8.2% |  |  |  | 1.334111 | - 75.817854 |
| 15 | M_017 | 7.5% |  |  |  | 1.337160 | - 75.808508 |
| 16 | M_018 |  |  |  | 45.8% | 1.324906 | - 75.806795 |
| 17 | M_019 |  |  | 21.1% |  | 1.299224 | - 75.811378 |
| 18 | M_020 |  | 10.9% |  |  | 1.279758 | - 75.808022 |
| 19 | M_021 | 7.7% |  |  |  | 1.256783 | - 75.796746 |
| 20 | M_022 | 7.3% |  |  |  | 1.253573 | - 75.779521 |
| 21 | M_023 |  |  | 20.0% |  | 1.228224 | - 75.780593 |
| 22 | M_024 |  | 10.5% |  |  | 1.225511 | - 75.764755 |
| 23 | M_025 |  | 13.5% |  |  | 1.197095 | - 75.749842 |
| 24 | M_026 | 6.0% |  |  |  | 1.408262 | - 75.728417 |
| 25 | M_027 | 9.9% |  |  |  | 1.191588 | - 75.743620 |
| 26 | M_028 |  |  |  | 44.9% | 1.557929 | - 75.713066 |
| 27 | M_029 |  | 15.1% |  |  | 1.521835 | - 75.703549 |
| 28 | M_030 | 5.7% |  |  |  | 1.442468 | - 75.701566 |
| 29 | M_031 |  | 13.7% |  |  | 1.354132 | - 75.701106 |
| 30 | M_032 | 9.0% |  |  |  | 1.238044 | - 75.702889 |
| 31 | M_033 |  |  | 20.4% |  | 1.490218 | - 75.683298 |
| 32 | M_034 |  | 14.9% |  |  | 1.450677 | - 75.689406 |
| 33 | M_035 | 9.7% |  |  |  | 1.443084 | - 75.694352 |
| 34 | M_036 |  | 15.7% |  |  | 1.467633 | - 75.669171 |
| 35 | M_037 | 5.4% |  |  |  | 1.291609 | - 75.659198 |
| 36 | M_038 | 9.2% |  |  |  | 1.194058 | - 75.670468 |
| 37 | M_039 |  |  |  | 42.7% | 1.740201 | - 75.634167 |
| 38 | M_041 |  |  |  | 50.4% | 1.672856 | - 75.600615 |
| 39 | M_042 | 9.0% |  |  |  | 1.542241 | - 75.598059 |
| 40 | M_043 |  | 12.8% |  |  | 1.528427 | - 75.597234 |
| 41 | M_044 | 3.9% |  |  |  | 1.499354 | - 75.604701 |
| 42 | M_045 |  | 11.2% |  |  | 1.483424 | - 75.605059 |
| 43 | M_047 |  |  |  | 50.0% | 1.616939 | - 75.568907 |
| 44 | M_048 |  | 12.1% |  |  | 1.451918 | - 75.577694 |
| 45 | M_049 |  | 11.8% |  |  | 1.612318 | - 75.564270 |
| 46 | M_050 | 9.2% |  |  |  | 1.517332 | - 75.547405 |
| 47 | M_051 |  | 12.4% |  |  | 1.507634 | - 75.554480 |
| 48 | M_052 | 9.3% |  |  |  | 1.472857 | - 75.547012 |
| 49 | M_053 | 4.1% |  |  |  | 1.386701 | - 75.547708 |
| 50 | M_054 | 9.5% |  |  |  | 1.291460 | - 75.546079 |
| 51 | M_055 | 8.2% |  |  |  | 1.502155 | - 75.529685 |
| 52 | M_056 |  |  | 19.6% |  | 1.483046 | - 75.541733 |
| 53 | M_057 |  | 14.9% |  |  | 1.452263 | - 75.529762 |
| 54 | M_058 |  |  | 21.2% |  | 1.430541 | - 75.528456 |
| 55 | M_059 | 7.7% |  |  |  | 1.418228 | - 75.535257 |
| 56 | M_060 | 8.5% |  |  |  | 1.390002 | - 75.542616 |
| 57 | M_061 |  | 13.1% |  |  | 1.299633 | - 75.522655 |
| 58 | M_062 | 7.5% |  |  |  | 1.431397 | - 75.514396 |
| 59 | M_063 |  | 10.5% |  |  | 1.424759 | - 75.505496 |
| 60 | M_064 |  |  | 18.6% |  | 1.298879 | - 75.518998 |
| 61 | M_065 |  | 15.9% |  |  | 1.506100 | - 75.489884 |
| 62 | M_066 |  | 13.8% |  |  | 1.430745 | - 75.491599 |
| 63 | M_067 |  |  |  | 46.1% | 1.675576 | - 75.464258 |
| 64 | M_068 |  | 13.9% |  |  | 1.497228 | - 75.431594 |
| 65 | M_069 | 7.5% |  |  |  | 1.488325 | - 75.438956 |
| 66 | M_071 |  |  |  | 51.7% | 1.514707 | - 75.396168 |
| 67 | M_072 | 8.8% |  |  |  | 1.298391 | - 75.394028 |
| 68 | M_073 |  | 14.0% |  |  | 1.402602 | - 75.375602 |
| 69 | M_074 |  | 14.3% |  |  | 1.337392 | - 75.354971 |
| 70 | M_075 |  | 14.1% |  |  | 1.332364 | - 75.353899 |
| 71 | M_076 |  |  | 19.0% |  | 1.293929 | - 75.330456 |
| 72 | M_077 |  |  |  | 48.1% | 1.653133 | - 75.317524 |
| 73 | M_078 |  | 10.5% |  |  | 1.504209 | - 75.310445 |
| 74 | M_080 | 9.0% |  |  |  | 1.282652 | - 75.301244 |
| 75 | M_081 |  | 13.1% |  |  | 1.295616 | - 75.291146 |
| 76 | M_082 |  |  |  | 74.0% | 1.838399 | - 75.257885 |
| 77 | M_083 | 8.4% |  |  |  | 1.742832 | - 75.263892 |
| 78 | M_084 | 7.5% |  |  |  | 1.738159 | - 75.257766 |
| 79 | M_085 |  |  | 19.3% |  | 1.643669 | - 75.262406 |
| 80 | M_086 | 6.3% |  |  |  | 1.752983 | - 75.245570 |
| 81 | M_087 |  |  | 19.7% |  | 1.733711 | - 75.249572 |
| 82 | M_088 |  |  | 21.7% |  | 1.759206 | - 75.217329 |
| 83 | M_089 |  |  | 20.3% |  | 1.848017 | - 75.187263 |
| 84 | M_090 | 9.1% |  |  |  | 1.601238 | - 75.204671 |
| 85 | M_092 |  |  | 20.9% |  | 1.648803 | - 75.122035 |
| 86 | M_093 |  | 12.0% |  |  | 2.006500 | - 75.025650 |
| 87 | M_094 | 5.3% |  |  |  | 1.784924 | - 74.998425 |
| 88 | M_095 |  | 11.3% |  |  | 1.630041 | - 74.993879 |
| 89 | M_096 |  | 11.0% |  |  | 1.925147 | - 74.980603 |
| 90 | M_097 |  |  | 20.9% |  | 2.105772 | - 74.830812 |
| 91 | M_098 | 5.0% |  |  |  | 1.990896 | - 74.834580 |
| 92 | M_099 |  |  |  | 42.2% | 2.110209 | - 74.580074 |
| 93 | M_100 |  |  |  | 48.0% | 2.103302 | - 74.577938 |
| 94 | M_101 | 4.5% |  |  |  | 2.108990 | - 74.566139 |
| 95 | M_102 |  | 10.9% |  |  | 2.153142 | - 74.545279 |
| 96 | M_103 |  | 13.4% |  |  | 2.148326 | - 74.546440 |
| 97 | M_104 | 8.2% |  |  |  | 2.108399 | - 74.553748 |
| 98 | M_105 |  |  | 21.3% |  | 2.143670 | - 74.519893 |
| 99 | M_106 |  | 11.0% |  |  | 1.814057 | - 74.513853 |
| 100 | M_107 |  | 11.0% |  |  | 2.049642 | - 74.353396 |
| 101 | M_108 |  |  |  | 42.8% | 1.869016 | - 74.241061 |
| 102 | M_109 |  |  | 22.6% |  | 2.168043 | - 74.056822 |
| 103 | M_110 | 7.9% |  |  |  | 1.174105 | - 75.330351 |
| 104 | M_111 |  |  | 19.2% |  | 1.160347 | - 75.253770 |

OP: open; SO: semi-open; SC: semi-closed; CL: closed.
