## Supplementary material for "Tree-Cover Gradients Modulate the Taxonomic and Functional Diversity of Birds in Amazonian Cattle-Ranching Landscapes": Table S2

**Table S2.** Bird species composition and guild assignment across tree cover gradients in Amazonian ranching landscapes.

| **Order** | **Family** | **Scientific name** | **Guild** | **CL** | **OP** | **SC** | **SO** | **Total** |
| --- | --- | --- | --- | --- | --- | --- | --- | --- |
| Accipitriformes | Accipitridae | *Buteo brachyurus* | VER |  | 3 |  |  | 3 |
|  |  | *Buteo nitidus* | VER | 9 | 11 | 1 | 9 | 30 |
|  |  | *Buteogallus meridionalis* | VER |  | 3 | 1 |  | 4 |
|  |  | *Chondrohierax uncinatus* | VER | 1 | 1 |  | 2 | 4 |
|  |  | *Elanoides forficatus* | VER |  | 2 | 29 | 25 | 56 |
|  |  | *Elanus leucurus* | VER |  | 3 | 1 |  | 4 |
|  |  | *Gampsonyx swainsonii* | VER |  |  |  | 1 | 1 |
|  |  | *Geranospiza caerulescens* | VER |  | 1 |  | 2 | 3 |
|  |  | *Leucopternis semiplumbeus* | VER |  | 1 |  | 1 | 2 |
|  |  | *Rupornis magnirostris* | VER | 20 | 63 | 38 | 86 | 207 |
|  |  | *Spizaetus tyrannus* | VER |  | 1 |  |  | 1 |
|  | Pandionidae | *Pandion haliaetus* | VER |  | 1 | 2 |  | 3 |
| Anseriformes | Anatidae | *Dendrocygna autumnalis* | INS | 2 | 19 | 5 | 5 | 31 |
| Apodiformes | Apodidae | *Chaetura brachyura* | INS |  | 1 |  |  | 1 |
|  |  | *Chaetura cinereiventris* | INS | 1 | 21 | 5 | 14 | 41 |
|  |  | *Cypseloides cryptus* | INS | 1 | 60 |  | 23 | 84 |
|  |  | *Streptoprocne zonaris* | INS | 1 | 86 | 10 | 9 | 106 |
|  |  | *Tachornis squamata* | INS | 1 | 19 |  | 25 | 45 |
| Caprimulgiformes | Caprimulgidae | *Chordeiles minor* | INS |  | 1 |  |  | 1 |
|  |  | *Nyctidromus albicollis* | INS | 1 | 3 |  | 1 | 5 |
| Charadriiformes | Charadriidae | *Hoploxypterus cayanus* | INS |  | 2 |  |  | 2 |
|  |  | *Vanellus chilensis* | INS | 22 | 154 | 55 | 148 | 379 |
|  | Jacanidae | *Jacana jacana* | INS | 6 | 30 | 24 | 35 | 95 |
|  | Rynchopidae | *Rynchops niger* | INS |  | 2 |  |  | 2 |
|  | Scolopacidae | *Actitis macularius* | INS |  | 1 |  | 1 | 2 |
|  |  | *Calidris minutilla* | INS |  | 1 |  |  | 1 |
|  |  | *Tringa solitaria* | INS | 2 | 2 | 3 |  | 7 |
| Columbiformes | Columbidae | *Columbina minuta* | GRA | 4 | 16 | 4 | 17 | 41 |
|  |  | *Columbina squammata* | GRA |  | 3 | 9 | 2 | 14 |
|  |  | *Columbina talpacoti* | GRA | 18 | 69 | 30 | 80 | 197 |
|  |  | *Geotrygon montana* | GRA |  | 2 |  |  | 2 |
|  |  | *Leptotila rufaxilla* | GRA | 7 | 19 | 15 | 18 | 59 |
|  |  | *Leptotila verreauxi* | GRA |  | 1 |  |  | 1 |
|  |  | *Patagioenas cayennensis* | GRA | 26 | 91 | 66 | 86 | 269 |
|  |  | *Patagioenas plumbea* | GRA | 2 | 24 | 5 | 19 | 50 |
|  |  | *Patagioenas speciosa* | GRA | 1 | 2 | 1 | 1 | 5 |
|  |  | *Patagioenas subvinacea* | GRA |  | 4 | 1 | 7 | 12 |
|  |  | *Zenaida auriculata* | GRA |  |  | 2 |  | 2 |
| Coraciiformes | Alcedinidae | *Chloroceryle aenea* | VER |  |  |  | 2 | 2 |
|  |  | *Chloroceryle amazona* | VER |  | 5 | 3 | 7 | 15 |
|  |  | *Chloroceryle americana* | VER | 2 | 3 | 2 | 3 | 10 |
|  |  | *Megaceryle torquata* | VER | 4 | 8 | 1 | 4 | 17 |
|  | Momotidae | *Baryphthengus martii* | INS | 2 |  | 1 | 1 | 4 |
|  |  | *Momotus momota* | INS | 1 |  | 4 | 1 | 6 |
| Cuculiformes | Cuculidae | *Coccycua minuta* | INS |  | 6 | 3 | 2 | 11 |
|  |  | *Coccyzus americanus* | INS | 2 | 2 |  |  | 4 |
|  |  | *Coccyzus melacoryphus* | INS |  | 2 |  | 1 | 3 |
|  |  | *Crotophaga ani* | INS | 50 | 347 | 167 | 310 | 874 |
|  |  | *Crotophaga major* | INS | 2 | 49 | 34 | 58 | 143 |
|  |  | *Crotophaga sulcirostris* | INS |  | 4 |  | 3 | 7 |
|  |  | *Piaya cayana* | INS | 11 | 19 | 7 | 13 | 50 |
|  |  | *Tapera naevia* | INS |  | 2 |  | 2 | 4 |
| Eurypygiformes | Eurypygidae | *Eurypyga helias* | INS | 1 | 1 | 4 | 2 | 8 |
| Falconiformes | Falconidae | *Caracara cheriway* | VER |  |  |  | 1 | 1 |
|  |  | *Caracara plancus* | VER | 1 | 3 | 3 | 2 | 9 |
|  |  | *Daptrius ater* | VER |  | 23 | 16 | 34 | 73 |
|  |  | *Daptrius chimachima* | VER | 10 | 94 | 40 | 72 | 216 |
|  |  | *Falco rufigularis* | VER |  | 5 | 1 | 2 | 8 |
|  |  | *Falco sparverius* | VER |  | 2 |  | 4 | 6 |
|  |  | *Herpetotheres cachinnans* | VER |  | 5 | 4 | 1 | 10 |
|  |  | *Ibycter americanus* | VER |  | 2 | 5 | 13 | 20 |
| Galbuliformes | Bucconidae | *Chelidoptera tenebrosa* | INS | 2 |  |  | 2 | 4 |
|  |  | *Monasa flavirostris* | INS |  |  | 1 | 1 | 2 |
|  |  | *Monasa morphoeus* | INS | 1 |  |  |  | 1 |
|  |  | *Monasa nigrifrons* | INS | 3 | 14 | 5 | 18 | 40 |
|  | Galbulidae | *Brachygalba lugubris* | INS | 3 | 3 | 5 | 7 | 18 |
|  |  | *Galbalcyrhynchus leucotis* | INS | 2 | 12 | 3 | 8 | 25 |
|  |  | *Galbula dea* | INS |  | 1 |  |  | 1 |
|  |  | *Galbula galbula* | INS |  | 1 |  |  | 1 |
|  |  | *Galbula leucogastra* | INS | 1 |  |  |  | 1 |
|  |  | *Galbula tombacea* | INS | 1 | 2 | 5 | 13 | 21 |
| Galliformes | Cracidae | *Ortalis guttata* | FRU | 28 | 27 | 19 | 46 | 120 |
|  |  | *Ortalis motmot* | FRU | 21 | 19 | 6 | 15 | 61 |
|  |  | *Penelope jacquacu* | FRU | 2 | 3 | 4 | 2 | 11 |
|  |  | *Penelope perspicax* | FRU | 1 | 4 |  |  | 5 |
| Gruiformes | Heliornithidae | *Heliornis fulica* | INS | 1 |  |  |  | 1 |
|  | Rallidae | *Aramides cajaneus* | INS |  |  | 1 | 2 | 3 |
|  |  | *Laterallus exilis* | INS | 1 | 6 | 3 | 4 | 14 |
|  |  | *Laterallus melanophaius* | INS |  | 4 |  | 3 | 7 |
|  |  | *Porphyrio flavirostris* | INS |  |  |  | 2 | 2 |
|  |  | *Porphyrio martinica* | INS | 3 | 7 | 3 |  | 13 |
| Nyctibiiformes | Nyctibiidae | *Nyctibius grandis* | INS |  | 2 | 1 |  | 3 |
| Passeriformes | Cardinalidae | *Chlorothraupis carmioli* | INS |  |  |  | 1 | 1 |
|  |  | *Piranga flava* | FRU |  |  |  | 1 | 1 |
|  |  | *Piranga olivacea* | FRU | 2 |  |  |  | 2 |
|  |  | *Piranga rubra* | FRU | 2 | 9 |  | 4 | 15 |
|  | Corvidae | *Cyanocorax violaceus* | INS | 49 | 105 | 69 | 139 | 362 |
|  | Cotingidae | *Cephalopterus ornatus* | FRU |  | 2 |  |  | 2 |
|  |  | *Gymnoderus foetidus* | FRU |  | 1 |  |  | 1 |
|  |  | *Rupicola peruvianus* | FRU | 1 |  |  |  | 1 |
|  | Doncabobiidae | *Donacobius atricapilla* | INS |  | 29 | 14 | 15 | 58 |
|  | Fringillidae | *Euphonia chlorotica* | FRU |  |  | 2 | 2 | 4 |
|  |  | *Euphonia chrysopasta* | FRU | 3 | 8 | 8 | 14 | 33 |
|  |  | *Euphonia laniirostris* | FRU | 9 | 16 | 8 | 10 | 43 |
|  |  | *Euphonia mesochrysa* | FRU | 3 | 2 |  | 1 | 6 |
|  |  | *Euphonia minuta* | FRU |  |  |  | 2 | 2 |
|  |  | *Euphonia trinitatis* | FRU |  | 2 | 1 |  | 3 |
|  |  | *Euphonia xanthogaster* | FRU |  | 8 | 2 | 3 | 13 |
|  | Furnariidae | *Dendrocincla fuliginosa* | INS | 7 | 8 | 1 |  | 16 |
|  |  | *Dendrocincla merula* | INS |  | 1 |  |  | 1 |
|  |  | *Dendroplex kienerii* | INS |  |  |  | 1 | 1 |
|  |  | *Dendroplex picus* | INS | 7 | 31 | 11 | 28 | 77 |
|  |  | *Glyphorynchus spirurus* | INS |  | 1 |  |  | 1 |
|  |  | *Metopothrix aurantiaca* | INS |  | 20 | 1 | 13 | 34 |
|  |  | *Sittasomus griseicapillus* | INS |  |  |  | 1 | 1 |
|  |  | *Synallaxis albescens* | INS |  | 3 | 2 | 5 | 10 |
|  |  | *Synallaxis albigularis* | INS |  | 22 | 2 | 2 | 26 |
|  |  | *Xiphorhynchus guttatus* | INS |  | 1 |  |  | 1 |
|  | Hirundinidae | *Alopochelidon fucata* | INS | 2 |  |  |  | 2 |
|  |  | *Atticora fasciata* | INS | 4 | 44 | 9 | 1 | 58 |
|  |  | *Hirundo rustica* | INS |  |  | 3 |  | 3 |
|  |  | *Neochelidon tibialis* | INS | 1 | 1 | 1 | 3 | 6 |
|  |  | *Progne chalybea* | INS | 1 |  | 5 | 1 | 7 |
|  |  | *Progne modesta* | INS |  | 20 | 3 | 45 | 68 |
|  |  | *Progne tapera* | INS |  | 8 | 3 | 9 | 20 |
|  |  | *Pygochelidon cyanoleuca* | INS | 8 | 11 | 1 | 24 | 44 |
|  |  | *Stelgidopteryx ruficollis* | INS | 6 | 1 | 22 | 1 | 30 |
|  |  | *Tachycineta albiventer* | INS |  | 12 |  | 6 | 18 |
|  | Icteridae | *Cacicus cela* | FRU | 52 | 130 | 52 | 93 | 327 |
|  |  | *Cacicus latirostris* | FRU |  | 2 | 2 | 5 | 9 |
|  |  | *Cacicus solitarius* | FRU |  | 5 |  | 3 | 8 |
|  |  | *Gymnomystax mexicanus* | INS |  |  |  | 2 | 2 |
|  |  | *Icterus chrysater* | FRU |  | 1 |  |  | 1 |
|  |  | *Icterus chrysocephalus* | FRU |  | 2 |  |  | 2 |
|  |  | *Icterus croconotus* | FRU |  | 3 |  |  | 3 |
|  |  | *Leistes militaris* | INS | 1 | 44 | 44 | 45 | 134 |
|  |  | *Molothrus bonariensis* | INS | 2 | 10 | 2 | 7 | 21 |
|  |  | *Molothrus oryzivorus* | INS |  | 2 |  | 2 | 4 |
|  |  | *Psarocolius angustifrons* | FRU | 66 | 125 | 77 | 101 | 369 |
|  |  | *Psarocolius decumanus* | FRU | 33 | 102 | 48 | 94 | 277 |
|  |  | *Psarocolius viridis* | FRU |  |  |  | 1 | 1 |
|  |  | *Psarocolius wagleri* | FRU | 1 | 2 |  |  | 3 |
|  |  | *Sturnella magna* | INS | 4 | 18 | 21 | 28 | 71 |
|  | Mimidae | *Mimus gilvus* | INS | 2 | 12 | 7 | 22 | 43 |
|  | Parulidae | *Mniotilta varia* | INS |  | 1 |  |  | 1 |
|  |  | *Setophaga castanea* | INS |  | 5 |  |  | 5 |
|  |  | *Setophaga cerulea* | INS |  | 1 |  |  | 1 |
|  |  | *Setophaga fusca* | INS |  | 2 |  | 1 | 3 |
|  |  | *Setophaga petechia* | INS |  | 1 | 1 |  | 2 |
|  |  | *Setophaga ruticilla* | INS | 1 | 3 | 2 | 2 | 8 |
|  |  | *Setophaga striata* | INS | 6 | 6 | 9 | 1 | 22 |
|  | Passerellidae | *Ammodramus aurifrons* | GRA | 14 | 86 | 48 | 71 | 219 |
|  |  | *Arremonops conirostris* | GRA | 1 | 32 | 21 | 31 | 85 |
|  | Pipridae | *Lepidothrix coronata* | FRU |  |  | 4 | 1 | 5 |
|  |  | *Machaeropterus regulus* | FRU |  | 1 |  |  | 1 |
|  |  | *Machaeropterus striolatus* | FRU |  |  | 1 | 2 | 3 |
|  |  | *Manacus manacus* | FRU | 11 | 11 | 18 | 19 | 59 |
|  | Thamnophilidae | *Cercomacra nigricans* | INS |  | 1 | 1 |  | 2 |
|  |  | *Cercomacroides tyrannina* | INS | 4 |  |  |  | 4 |
|  |  | *Cymbilaimus lineatus* | INS |  |  | 1 |  | 1 |
|  |  | *Hypocnemis peruviana* | INS |  |  |  | 2 | 2 |
|  |  | *Myrmelastes leucostigma* | INS |  | 3 | 2 | 7 | 12 |
|  |  | *Myrmotherula axillaris* | INS | 1 | 2 |  |  | 3 |
|  |  | *Myrmotherula brachyura* | INS |  |  | 2 |  | 2 |
|  |  | *Thamnophilus amazonicus* | INS |  |  | 2 |  | 2 |
|  |  | *Thamnophilus schistaceus* | INS |  |  | 2 |  | 2 |
|  | Thraupidae | *Cissopis leverianus* | FRU | 6 | 14 | 7 | 6 | 33 |
|  |  | *Dacnis berlepschi* | FRU |  |  |  | 2 | 2 |
|  |  | *Dacnis cayana* | FRU | 2 |  |  |  | 2 |
|  |  | *Ixothraupis xanthogastra* | FRU | 7 |  |  |  | 7 |
|  |  | *Loriotus cristatus* | FRU |  | 1 |  |  | 1 |
|  |  | *Paroaria gularis* | FRU | 10 | 28 | 13 | 26 | 77 |
|  |  | *Ramphocelus carbo* | FRU | 52 | 156 | 98 | 170 | 476 |
|  |  | *Ramphocelus nigrogularis* | FRU | 1 | 5 |  | 5 | 11 |
|  |  | *Saltator coerulescens* | FRU | 3 | 13 | 11 | 8 | 35 |
|  |  | *Saltator maximus* | FRU | 9 | 11 | 11 | 24 | 55 |
|  |  | *Schistochlamys melanopis* | FRU | 6 | 8 | 12 | 16 | 42 |
|  |  | *Sicalis flaveola* | GRA | 6 | 60 | 32 | 61 | 159 |
|  |  | *Sporophila angolensis* | GRA | 10 | 18 | 10 | 22 | 60 |
|  |  | *Sporophila castaneiventris* | GRA |  | 4 |  | 5 | 9 |
|  |  | *Sporophila crassirostris* | GRA |  | 2 |  |  | 2 |
|  |  | *Sporophila intermedia* | GRA |  |  | 1 |  | 1 |
|  |  | *Sporophila minuta* | GRA | 3 |  | 1 |  | 4 |
|  |  | *Sporophila murallae* | GRA |  | 4 | 6 | 5 | 15 |
|  |  | *Sporophila nigricollis* | GRA |  |  |  | 2 | 2 |
|  |  | *Stilpnia nigrocincta* | FRU | 2 |  | 11 | 9 | 22 |
|  |  | *Tachyphonus phoenicius* | FRU |  | 1 |  |  | 1 |
|  |  | *Tachyphonus rufus* | FRU | 2 | 11 | 2 | 8 | 23 |
|  |  | *Tachyphonus surinamus* | FRU | 11 | 2 |  |  | 13 |
|  |  | *Tangara chilensis* | FRU |  |  | 3 |  | 3 |
|  |  | *Tangara mexicana* | FRU | 10 | 26 | 25 | 16 | 77 |
|  |  | *Tangara velia* | FRU | 1 |  |  |  | 1 |
|  |  | *Tersina viridis* | FRU | 8 | 10 | 23 | 2 | 43 |
|  |  | *Thraupis episcopus* | FRU | 61 | 237 | 113 | 209 | 620 |
|  |  | *Thraupis palmarum* | FRU | 36 | 149 | 52 | 148 | 385 |
|  |  | *Volatinia jacarina* | GRA | 11 | 72 | 43 | 61 | 187 |
|  | Tityridae | *Pachyramphus albogriseus* | INS |  |  | 6 |  | 6 |
|  |  | *Pachyramphus marginatus* | INS | 2 | 1 | 3 |  | 6 |
|  |  | *Pachyramphus polychopterus* | INS | 4 | 24 | 25 | 31 | 84 |
|  |  | *Pachyramphus rufus* | INS |  |  | 1 | 11 | 12 |
|  |  | *Tityra cayana* | INS | 8 | 21 | 12 | 24 | 65 |
|  |  | *Tityra inquisitor* | INS | 4 | 15 | 1 | 2 | 22 |
|  |  | *Tityra semifasciata* | INS | 2 | 3 | 1 | 16 | 22 |
|  | Troglodytidae | *Campylorhynchus turdinus* | INS |  | 3 |  | 2 | 5 |
|  |  | *Cantorchilus leucotis* | INS |  | 8 | 11 | 7 | 26 |
|  |  | *Henicorhina leucosticta* | INS | 2 |  | 1 |  | 3 |
|  |  | *Microcerculus marginatus* | INS |  | 2 |  |  | 2 |
|  |  | *Pheugopedius coraya* | INS |  |  | 2 | 4 | 6 |
|  |  | *Troglodytes aedon* | INS | 19 | 59 | 19 | 85 | 182 |
|  | Turdidae | *Catharus ustulatus* | INS | 2 |  |  |  | 2 |
|  |  | *Turdus ignobilis* | INS | 13 | 67 | 9 | 26 | 115 |
|  | Tyrannidae | *Attila cinnamomeus* | INS |  |  | 2 |  | 2 |
|  |  | *Attila citriniventris* | INS |  |  |  | 1 | 1 |
|  |  | *Attila spadiceus* | INS | 2 | 9 | 4 | 9 | 24 |
|  |  | *Camptostoma obsoletum* | INS |  | 16 | 14 | 16 | 46 |
|  |  | *Capsiempis flaveola* | INS |  |  | 2 |  | 2 |
|  |  | *Conopias parvus* | INS |  | 5 | 3 | 6 | 14 |
|  |  | *Contopus cinereus* | INS |  | 2 |  |  | 2 |
|  |  | *Contopus virens* | INS |  | 3 | 4 |  | 7 |
|  |  | *Elaenia flavogaster* | INS | 2 | 10 |  | 9 | 21 |
|  |  | *Elaenia frantzii* | FRU |  |  |  | 3 | 3 |
|  |  | *Elaenia parvirostris* | INS |  |  |  | 3 | 3 |
|  |  | *Empidonax alnorum* | INS | 2 |  |  |  | 2 |
|  |  | *Empidonomus varius* | INS | 1 | 17 | 2 | 12 | 32 |
|  |  | *Hemitriccus striaticollis* | INS | 1 |  |  |  | 1 |
|  |  | *Hemitriccus zosterops* | INS | 5 | 27 | 13 | 19 | 64 |
|  |  | *Lathrotriccus euleri* | INS |  | 2 |  | 2 | 4 |
|  |  | *Legatus leucophaius* | INS | 1 | 8 | 5 | 18 | 32 |
|  |  | *Leptopogon amaurocephalus* | INS | 1 |  | 4 | 3 | 8 |
|  |  | *Machetornis rixosa* | INS |  |  |  | 2 | 2 |
|  |  | *Megarynchus pitangua* | INS | 10 | 28 | 22 | 29 | 89 |
|  |  | *Mionectes oleagineus* | INS |  | 2 |  | 2 | 4 |
|  |  | *Myiarchus crinitus* | INS | 1 |  |  |  | 1 |
|  |  | *Myiarchus ferox* | INS | 7 | 23 | 44 | 24 | 98 |
|  |  | *Myiarchus swainsoni* | INS |  | 1 |  |  | 1 |
|  |  | *Myiarchus tuberculifer* | INS |  | 10 | 11 | 6 | 27 |
|  |  | *Myiarchus venezuelensis* | INS |  | 1 |  | 1 | 2 |
|  |  | *Myiodynastes maculatus* | INS | 8 | 19 | 12 | 7 | 46 |
|  |  | *Myiopagis caniceps* | INS | 1 | 1 |  |  | 2 |
|  |  | *Myiopagis gaimardii* | INS | 2 |  |  |  | 2 |
|  |  | *Myiophobus flavicans* | INS |  | 2 |  |  | 2 |
|  |  | *Myiozetetes cayanensis* | INS | 3 | 22 | 8 | 2 | 35 |
|  |  | *Myiozetetes granadensis* | INS |  | 14 | 4 | 14 | 32 |
|  |  | *Myiozetetes luteiventris* | INS |  |  |  | 4 | 4 |
|  |  | *Myiozetetes similis* | INS | 22 | 101 | 52 | 65 | 240 |
|  |  | *Nesotriccus murinus* | INS | 2 | 5 |  | 1 | 8 |
|  |  | *Philohydor lictor* | INS | 2 | 10 | 6 | 14 | 32 |
|  |  | *Pitangus sulphuratus* | INS | 29 | 110 | 54 | 125 | 318 |
|  |  | *Poecilotriccus latirostris* | INS |  | 1 |  | 1 | 2 |
|  |  | *Pyrocephalus rubinus* | INS |  | 4 | 4 | 14 | 22 |
|  |  | *Rhynchocyclus olivaceus* | INS |  | 2 |  |  | 2 |
|  |  | *Sayornis nigricans* | INS | 10 | 1 | 3 |  | 14 |
|  |  | *Sirystes sibilator* | INS | 1 | 1 | 2 | 1 | 5 |
|  |  | *Terenotriccus erythrurus* | INS |  |  | 3 | 2 | 5 |
|  |  | *Todirostrum chrysocrotaphu* | INS |  | 17 | 14 | 15 | 46 |
|  |  | *Todirostrum cinereum* | INS | 35 | 54 | 41 | 52 | 182 |
|  |  | *Todirostrum maculatum* | INS |  | 6 | 3 | 1 | 10 |
|  |  | *Tolmomyias assimilis* | INS |  | 1 |  |  | 1 |
|  |  | *Tolmomyias poliocephalus* | INS |  | 9 | 7 | 3 | 19 |
|  |  | *Tolmomyias sulphurescens* | INS | 2 |  |  | 2 | 4 |
|  |  | *Tyrannopsis sulphurea* | INS |  | 3 |  | 6 | 9 |
|  |  | *Tyrannulus elatus* | INS | 2 | 19 | 4 | 12 | 37 |
|  |  | *Tyrannus melancholicus* | INS | 48 | 218 | 117 | 210 | 593 |
|  |  | *Tyrannus niveigularis* | INS |  | 1 |  |  | 1 |
|  |  | *Tyrannus savana* | INS | 4 | 53 | 27 | 36 | 120 |
|  |  | *Tyrannus tyrannus* | INS |  | 9 | 9 | 6 | 24 |
|  |  | *Zimmerius viridiflavus* | INS |  |  |  | 2 | 2 |
|  |  | *Zimmerius gracilipes* | INS |  | 1 |  |  | 1 |
|  | Vireonidae | *Cyclarhis gujanensis* | INS |  | 1 |  |  | 1 |
|  |  | *Vireo flavoviridis* | INS |  |  | 2 |  | 2 |
|  |  | *Vireo olivaceus* | FRU |  | 2 | 3 |  | 5 |
| Pelecaniformes | Ardeidae | *Ardea alba* | VER |  | 7 | 5 | 7 | 19 |
|  |  | *Ardea cocoi* | VER |  | 1 | 1 | 4 | 6 |
|  |  | *Ardea ibis* | INS | 102 | 423 | 158 | 462 | 1145 |
|  |  | *Butorides striata* | VER | 4 | 9 | 5 | 6 | 24 |
|  |  | *Egretta caerulea* | VER |  |  | 2 |  | 2 |
|  |  | *Egretta thula* | VER |  | 1 | 7 | 13 | 21 |
|  |  | *Pilherodius pileatus* | VER | 1 |  |  | 1 | 2 |
|  |  | *Syrigma sibilatrix* | INS | 5 | 13 | 9 | 17 | 44 |
|  |  | *Tigrisoma fasciatum* | VER | 2 | 10 | 2 | 2 | 16 |
|  |  | *Tigrisoma lineatum* | VER | 1 | 11 | 6 | 4 | 22 |
|  | Threskiornithidae | *Eudocimus ruber* | INS |  | 67 | 4 | 20 | 91 |
|  |  | *Mesembrinibis cayennensis* | INS | 7 | 26 | 11 | 23 | 67 |
|  |  | *Phimosus infuscatus* | INS | 18 | 80 | 22 | 80 | 200 |
| Piciformes | Capitonidae | *Capito auratus* | FRU | 3 | 9 | 28 | 24 | 64 |
|  |  | *Capito aurovirens* | FRU | 5 | 18 | 11 | 21 | 55 |
|  |  | *Capito niger* | FRU | 2 | 7 | 1 | 1 | 11 |
|  | Picidae | *Campephilus melanoleucos* | INS | 4 | 8 | 6 | 3 | 21 |
|  |  | *Celeus elegans* | INS |  | 2 |  | 8 | 10 |
|  |  | *Celeus flavus* | INS |  | 7 |  | 5 | 12 |
|  |  | *Colaptes punctigula* | INS | 1 | 15 | 1 | 10 | 27 |
|  |  | *Colaptes rubiginosus* | INS | 5 | 12 | 1 | 4 | 22 |
|  |  | *Dryobates affinis* | INS | 1 | 2 | 1 |  | 4 |
|  |  | *Dryobates passerinus* | INS |  |  |  | 3 | 3 |
|  |  | *Dryocopus lineatus* | INS |  | 13 | 6 | 15 | 34 |
|  |  | *Melanerpes cruentatus* | INS | 17 | 70 | 33 | 63 | 183 |
|  |  | *Picoides fumigatus* | INS |  |  | 1 |  | 1 |
|  |  | *Picumnus lafresnayi* | INS | 6 | 13 | 8 | 12 | 39 |
|  |  | *Picumnus squamulatus* | INS | 1 | 3 |  |  | 4 |
|  | Ramphastidae | *Pteroglossus castanotis* | FRU | 5 | 12 | 14 | 15 | 46 |
|  |  | *Pteroglossus inscriptus* | FRU | 6 | 29 | 3 | 16 | 54 |
|  |  | *Pteroglossus pluricinctus* | FRU | 8 | 18 | 6 | 15 | 47 |
|  |  | *Pteroglossus torquatus* | FRU |  | 2 |  |  | 2 |
|  |  | *Ramphastos tucanus* | FRU | 4 | 4 | 12 | 13 | 33 |
| Podicipediformes | Podicipedidae | *Tachybaptus dominicus* | INS |  |  | 1 | 3 | 4 |
| Psittaciformes | Psittacidae | *Amazona aestiva* | FRU |  | 2 | 2 |  | 4 |
|  |  | *Amazona amazonica* | FRU | 4 | 100 | 28 | 71 | 203 |
|  |  | *Amazona farinosa* | FRU |  | 11 | 3 | 3 | 17 |
|  |  | *Amazona ochrocephala* | FRU | 2 | 108 | 110 | 122 | 342 |
|  |  | *Ara ararauna* | FRU | 1 | 10 | 10 | 12 | 33 |
|  |  | *Ara macao* | FRU |  | 8 | 5 | 9 | 22 |
|  |  | *Ara severus* | FRU | 97 | 295 | 138 | 392 | 922 |
|  |  | *Aratinga weddellii* | FRU | 2 | 24 | 9 | 59 | 94 |
|  |  | *Brotogeris cyanoptera* | FRU | 23 | 94 | 52 | 150 | 319 |
|  |  | *Eupsittula pertinax* | FRU | 2 |  | 4 | 3 | 9 |
|  |  | *Forpus conspicillatus* | FRU |  |  | 1 |  | 1 |
|  |  | *Forpus crassirostris* | FRU |  |  |  | 5 | 5 |
|  |  | *Forpus modestus* | FRU |  | 10 |  | 4 | 14 |
|  |  | *Orthopsittaca manilatus* | FRU | 2 | 24 |  |  | 26 |
|  |  | *Pionites melanocephalus* | FRU | 16 | 18 | 7 | 13 | 54 |
|  |  | *Pionus menstruus* | FRU | 2 | 55 | 22 | 54 | 133 |
|  |  | *Psittacara leucophthalmus* | FRU | 6 | 16 | 3 | 9 | 34 |
|  |  | *Pyrilia barrabandi* | FRU |  |  | 5 |  | 5 |
| Strigiformes | Strigidae | *Glaucidium brasilianum* | VER | 1 | 2 | 2 |  | 5 |
|  |  | *Megascops choliba* | VER |  | 4 |  | 1 | 5 |
|  | Tytonidae | *Tyto alba* | VER |  |  |  | 1 | 1 |
| Suliformes | Phalacrocoracidae | *Nannopterum brasilianum* | VER |  | 1 | 2 |  | 3 |
| Tinamiformes | Tinamidae | *Crypturellus cinereus* | GRA | 4 | 4 | 1 | 2 | 11 |
|  |  | *Crypturellus soui* | GRA | 1 | 1 | 4 | 7 | 13 |
|  |  | *Crypturellus undulatus* | GRA | 1 | 5 | 4 | 6 | 16 |
|  |  | *Tinamus guttatus* | GRA | 1 |  | 3 |  | 4 |
|  |  | *Tinamus tao* | GRA |  | 1 |  |  | 1 |
| Trogoniformes | Trogonidae | *Trogon collaris* | INS | 2 |  |  |  | 2 |
|  |  | *Trogon viridis* | INS | 5 | 17 | 14 | 27 | 63 |
| **Total** | | |  | **1482** | **5977** | **3057** | **5743** | **16259** |

SO: semi-open; OP: open; SC: semi-closed; CL: closed; VER: vertebrate consumers; GRA: granivorous; FRU: frugivorous; INS: insectivorous.
