## Supplementary material S3 for "Tree-Cover Gradients Modulate the Taxonomic and Functional Diversity of Birds in Amazonian Cattle-Ranching Landscapes"

**Supplementary material S3.** Mixed Linear Models taxonomic diversity and Functional diversity metrics across vegetation mosaics the livestock landscapes of northwestern Amazonia in Colombia

**Open vegetation mosaics (OP)**

***Variable = Richness***

| **Fountain** | **GLNum** | **GLDen** | **F** | **P-Value** |
| --- | --- | --- | --- | --- |
| (Intercept) | 1 | 137 | 246.46 | <0.0001 |
| Guild | 3 | 137 | 27.06 | <0.0001 |

*Average Comparisons (Guild)*

Method of comparing pairs of means: Fisher. alpha=0.05

Mean Difference Mean Standard Error=1.91

Minimum Average Significant Difference = 3.78

| **coverage** | **Variable** | **Guild** | **Stocking** | **USA** | **N** | **Group** |
| --- | --- | --- | --- | --- | --- | --- |
| open | Richness | INS | 19.93 | 1.19 | 44 | A |
| open | Richness | FRU | 10.79 | 1.28 | 38 | B |
| open | Richness | GRA | 5.85 | 1.52 | 27 | C |
| open | Richness | SEE | 5.81 | 1.39 | 32 | C |

Stockings with a common letter are not significantly different (e > 0.05).

***Variable = Abundance***

| **Fountain** | **GLNum** | **GLDen** | **F** | **P-Value** |
| --- | --- | --- | --- | --- |
| (Intercept) | 1 | 137 | 95.77 | <0.0001 |
| Guild | 3 | 137 | 7.66 | <0.0001 |

*Average Comparisons (Guild)*

Method of comparing pairs of means: Fisher. alpha=0.05

Mean difference average standard error=12.5

Minimum Average Significant Difference = 24.7

| **coverage** | **Variable** | **Guild** | **Stocking** | **USA** | **N** | **Group** |
| --- | --- | --- | --- | --- | --- | --- |
| open | Abundance | INS | 72.48 | 7.79 | 44 | A |
| open | Abundance | FRU | 52.82 | 8.38 | 38 | A |
| open | Abundance | SEE | 28.13 | 9.13 | 32 | B |
| open | Abundance | GRA | 19.74 | 9.94 | 27 | B |

Stockings with a common letter are not significantly different (e > 0.05).

***Variable = Dominance_D***

| **Fountain** | **GLNum** | **GLDen** | **F** | **P-Value** |
| --- | --- | --- | --- | --- |
| (Intercept) | 1 | 137 | 279.06 | <0.0001 |
| Guild | 3 | 137 | 16.10 | <0.0001 |

*Average Comparisons (Guild)*

Method of comparing pairs of means: Fisher. alpha=0.05

Mean Difference Mean Standard Error=0.032

Minimum average significant difference = 0.063

| **Coverage** | **Variable** | **Guild** | **Stocking** | **USA** | **N** | **Group** |
| --- | --- | --- | --- | --- | --- | --- |
| open | Dominance_D | SEE | 0.31 | 0.02 | 32 | A |
| open | Dominance_D | GRA | 0.18 | 0.03 | 27 | B |
| open | Dominance_D | FRU | 0.16 | 0.02 | 38 | B |
| open | Dominance_D | INS | 0.10 | 0.02 | 44 | C |

Stockings with a common letter are not significantly different (e > 0.05).

***Variable = Shannon_H***

| **Fountain** | **GLNum** | **GLDen** | **F** | **P-Value** |
| --- | --- | --- | --- | --- |
| (Intercept) | 1 | 137 | 1843.09 | <0.0001 |
| Guild | 3 | 137 | 35.24 | <0.0001 |

*Average Comparisons (Guild)*

Method of comparing pairs of means: Fisher. alpha=0.05

Mean Standard Difference Average Error=0.131

Minimum Average Significant Difference = 0.259

| **Coverage** | **Variable** | **Guild** | **Stocking** | **USA** | **N** | **Group** |
| --- | --- | --- | --- | --- | --- | --- |
| open | Shannon_H | INS | 2.68 | 0.08 | 44 | A |
| open | Shannon_H | FRU | 2.09 | 0.09 | 38 | B |
| open | Shannon_H | GRA | 1.72 | 0.10 | 27 | C |
| open | Shannon_H | SEE | 1.48 | 0.10 | 32 | C |

Stockings with a common letter are not significantly different (e > 0.05).

***Variable = Margalef***

| **Fountain** | **GLNum** | **GLDen** | **F** | **P-Value** |
| --- | --- | --- | --- | --- |
| (Intercept) | 1 | 137 | 485.20 | <0.0001 |
| Guild | 3 | 137 | 36.10 | <0.0001 |

*Average Comparisons (Guild)*

Method of comparing pairs of means: Fisher. alpha=0.05

Mean Standard Difference Average Error = 0.333

Minimum Average Significant Difference = 0.658

| **Coverage** | **Variable** | **Guild** | **Stocking** | **USA** | **N** | **Group** |
| --- | --- | --- | --- | --- | --- | --- |
| open | Margalef | INS | 4.50 | 0.21 | 44 | A |
| open | Margalef | FRU | 2.50 | 0.22 | 38 | B |
| open | Margalef | GRA | 1.73 | 0.26 | 27 | C |
| open | Margalef | SEE | 1.64 | 0.24 | 32 | C |

Stockings with a common letter are not significantly different (e > 0.05).

***Variable = FRic***

| **Fountain** | **GLNum** | **GLDen** | **F** | **P-Value** |
| --- | --- | --- | --- | --- |
| (Intercept) | 1 | 137 | 240.30 | <0.0001 |
| Guild | 3 | 137 | 3.75 | 0.0126 |

*Average Comparisons (Guild)*

Method of comparing pairs of means: Fisher. alpha=0.05

Mean Standard Difference Average Error=0.042

Minimum average significant difference = 0.083

| **Coverage** | **Variable** | **Guild** | **Stocking** | **USA** | **N** | **Group** |
| --- | --- | --- | --- | --- | --- | --- |
| open | FRic | INS | 0.30 | 0.03 | 44 | A |
| open | FRic | FRU | 0.24 | 0.03 | 38 | AB |
| open | FRic | GRA | 0.21 | 0.03 | 27 | B |
| open | FRic | SEE | 0.17 | 0.03 | 32 | B |

Stockings with a common letter are not significantly different (e > 0.05).

***Variable = FEve***

| **Fountain** | **GLNum** | **GLDen** | **F** | **P-Value** |
| --- | --- | --- | --- | --- |
| (Intercept) | 1 | 137 | 3228.24 | <0.0001 |
| Guild | 3 | 137 | 4.30 | 0.0062 |

*Average Comparisons (Guild)*

Method of comparing pairs of means: Fisher. alpha=0.05

Mean difference average standard error=0.034

Minimum average significant difference = 0.066

| **Coverage** | **Variable** | **Guild** | **Stocking** | **USA** | **N** | **Group** |
| --- | --- | --- | --- | --- | --- | --- |
| open | FAITH | INS | 0.71 | 0.02 | 44 | A |
| open | FAITH | GRA | 0.70 | 0.03 | 27 | A |
| open | FAITH | SEE | 0.69 | 0.02 | 32 | A |
| open | FAITH | FRU | 0.61 | 0.02 | 38 | B |

Stockings with a common letter are not significantly different (e > 0.05).

***Variable = FDiv***

| **Fountain** | **GLNum** | **GLDen** | **F** | **P-Value** |
| --- | --- | --- | --- | --- |
| (Intercept) | 1 | 137 | 6914.58 | <0.0001 |
| Guild | 3 | 137 | 10.75 | <0.0001 |

*Average Comparisons (Guild)*

Method of comparing pairs of means: Fisher. alpha=0.05

Mean difference average standard error=0.024

Minimum average significant difference = 0.048

| **Coverage** | **Variable** | **Guild** | **Stocking** | **USA** | **N** | **Group** |
| --- | --- | --- | --- | --- | --- | --- |
| open | FDiv | FRU | 0.77 | 0.02 | 38 | A |
| open | FDiv | GRA | 0.73 | 0.02 | 27 | A |
| open | FDiv | INS | 0.68 | 0.01 | 44 | B |
| open | FDiv | SEE | 0.65 | 0.02 | 32 | B |

Stockings with a common letter are not significantly different (e > 0.05).

***Variable = FDis***

| **Fountain** | **GLNum** | **GLDen** | **F** | **P-Value** |
| --- | --- | --- | --- | --- |
| (Intercept) | 1 | 137 | 1621.14 | <0.0001 |
| Guild | 3 | 137 | 44.62 | <0.0001 |

*Average Comparisons (Guild)*

Method of comparing pairs of means: Fisher. alpha=0.05

Mean Standard Difference Error = 0.161

Minimum average significant difference = 0.317

| **Coverage** | **Variable** | **Guild** | **Stocking** | **USA** | **N** | **Group** |
| --- | --- | --- | --- | --- | --- | --- |
| open | FDis | INS | 3.23 | 0.10 | 44 | A |
| open | FDis | FRU | 2.23 | 0.11 | 38 | B |
| open | FDis | GRA | 2.18 | 0.13 | 27 | B |
| open | FDis | SEE | 1.50 | 0.12 | 32 | C |

Stockings with a common letter are not significantly different (e > 0.05).

***Variable = RaoQ***

| **Fountain** | **GLNum** | **GLDen** | **F** | **P-Value** |
| --- | --- | --- | --- | --- |
| (Intercept) | 1 | 137 | 350.22 | <0.0001 |
| Guild | 3 | 137 | 39.22 | <0.0001 |

*Average Comparisons (Guild)*

Method of comparing pairs of means: Fisher. alpha=0.05

Mean difference average standard error=1.11

Minimum average significant difference = 2.2

| **Coverage** | **Variable** | **Guild** | **Stocking** | **USA** | **N** | **Group** |
| --- | --- | --- | --- | --- | --- | --- |
| open | RaoQ | INS | 13.96 | 0.69 | 44 | A |
| open | RaoQ | FRU | 6.34 | 0.75 | 38 | B |
| open | RaoQ | GRA | 5.89 | 0.89 | 27 | B |
| open | RaoQ | SEE | 3.31 | 0.81 | 32 | C |

Stockings with a common letter are not significantly different (e > 0.05).

**Mosaics semi-open vegetation (SO)**

***Variable = Richness***

| **Fountain** | **GLNum** | **GLDen** | **F** | **P-Value** |
| --- | --- | --- | --- | --- |
| (Intercept) | 1 | 43 | 101.30 | <0.0001 |
| Guild | 3 | 43 | 11.48 | <0.0001 |

*Average Comparisons (Guild)*

Method of comparing pairs of means: Fisher. alpha=0.05

Mean Difference Mean Standard Error=3.29

Minimum Average Significant Difference = 6.64

| **Coverage** | **Variable** | **Guild** | **Stocking** | **USA** | **N** | **Group** |
| --- | --- | --- | --- | --- | --- | --- |
| Semi-open | Richness | INS | 22.31 | 2.19 | 13 | A |
| Semi-open | Richness | FRU | 12.38 | 2.19 | 13 | B |
| Semi-open | Richness | GRA | 6.56 | 2.63 | 9 | BC |
| Semi-open | Richness | SEE | 5.58 | 2.28 | 12 | C |

Stockings with a common letter are not significantly different (e > 0.05).

***Variable = Abundance***

| **Fountain** | **GLNum** | **GLDen** | **F** | **P-Value** |
| --- | --- | --- | --- | --- |
| (Intercept) | 1 | 43 | 19.95 | <0.0001 |
| Guild | 3 | 43 | 2.04 | 0.1221 |

***Variable = Dominance_D***

| **Fountain** | **GLNum** | **GLDen** | **F** | **P-Value** |
| --- | --- | --- | --- | --- |
| (Intercept) | 1 | 43 | 133.48 | <0.0001 |
| Guild | 3 | 43 | 10.89 | <0.0001 |

*Average Comparisons (Guild)*

Method of comparing pairs of means: Fisher. alpha=0.05

Mean Difference Mean Standard Error=0.044

Minimum Average Significant Difference = 0.089

| **Coverage** | **Variable** | **Guild** | **Stocking** | **USA** | **N** | **Group** |
| --- | --- | --- | --- | --- | --- | --- |
| Semi-open | Dominance_D | SEE | 0.32 | 0.03 | 12 | A |
| Semi-open | Dominance_D | GRA | 0.17 | 0.04 | 9 | B |
| Semi-open | Dominance_D | FRU | 0.15 | 0.03 | 13 | B |
| Semi-open | Dominance_D | INS | 0.08 | 0.03 | 13 | B |

Stockings with a common letter are not significantly different (e > 0.05).

***Variable = Shannon_H***

| **Fountain** | **GLNum** | **GLDen** | **F** | **P-Value** |
| --- | --- | --- | --- | --- |
| (Intercept) | 1 | 43 | 925.56 | <0.0001 |
| Guild | 3 | 43 | 22.66 | <0.0001 |

*Average Comparisons (Guild)*

Method of comparing pairs of means: Fisher. alpha=0.05

Mean Standard Difference Average Error = 0.192

Minimum Average Significant Difference = 0.387

| **Coverage** | **Variable** | **Guild** | **Stocking** | **USA** | **N** | **Group** |
| --- | --- | --- | --- | --- | --- | --- |
| Semi-open | Shannon_H | INS | 2.88 | 0.13 | 13 | A |
| Semi-open | Shannon_H | FRU | 2.17 | 0.13 | 13 | B |
| Semi-open | Shannon_H | GRA | 1.78 | 0.15 | 9 | BC |
| Semi-open | Shannon_H | SEE | 1.42 | 0.13 | 12 | C |

Stockings with a common letter are not significantly different (e > 0.05).

***Variable = Margalef***

| **Fountain** | **GLNum** | **GLDen** | **F** | **P-Value** |
| --- | --- | --- | --- | --- |
| (Intercept) | 1 | 43 | 232.39 | <0.0001 |
| Guild | 3 | 43 | 18.71 | <0.0001 |

*Average Comparisons (Guild)*

Method of comparing pairs of means: Fisher. alpha=0.05

Mean Standard Difference Average Error=0.513

Minimum average significant difference = 1.03

| **Coverage** | **Variable** | **Guild** | **Stocking** | **USA** | **N** | **Group** |
| --- | --- | --- | --- | --- | --- | --- |
| Semi-open | Margalef | INS | 4.93 | 0.34 | 13 | A |
| Semi-open | Margalef | FRU | 2.68 | 0.34 | 13 | B |
| Semi-open | Margalef | GRA | 1.89 | 0.41 | 9 | BC |
| Semi-open | Margalef | SEE | 1.56 | 0.35 | 12 | C |

Stockings with a common letter are not significantly different (e > 0.05).

***Variable = FRic***

| **Fountain** | **GLNum** | **GLDen** | **F** | **P-Value** |
| --- | --- | --- | --- | --- |
| (Intercept) | 1 | 43 | 124.33 | <0.0001 |
| Guild | 3 | 43 | 1.53 | 0.2202 |

***Variable = FEve***

| **Fountain** | **GLNum** | **GLDen** | **F** | **P-Value** |
| --- | --- | --- | --- | --- |
| (Intercept) | 1 | 43 | 1526.26 | <0.0001 |
| Guild | 3 | 43 | 1.98 | 0.1318 |

***Variable = FDiv***

| **Fountain** | **GLNum** | **GLDen** | **F** | **P-Value** |
| --- | --- | --- | --- | --- |
| (Intercept) | 1 | 43 | 1775.04 | <0.0001 |
| Guild | 3 | 43 | 3.00 | 0.0406 |

*Average Comparisons (Guild)*

Method of comparing pairs of means: Fisher. alpha=0.05

Mean Difference Mean Standard Error=0.048

Minimum average significant difference = 0.098

| **Coverage** | **Variable** | **Guild** | **Stocking** | **USA** | **N** | **Group** |
| --- | --- | --- | --- | --- | --- | --- |
| Semi-open | FDiv | GRA | 0.78 | 0.04 | 9 | A |
| Semi-open | FDiv | FRU | 0.76 | 0.03 | 13 | A |
| Semi-open | FDiv | INS | 0.68 | 0.03 | 13 | AB |
| Semi-open | FDiv | SEE | 0.66 | 0.03 | 12 | B |

Stockings with a common letter are not significantly different (e > 0.05).

***Variable = FDis***

| **Fountain** | **GLNum** | **GLDen** | **F** | **P-Value** |
| --- | --- | --- | --- | --- |
| (Intercept) | 1 | 43 | 613.99 | <0.0001 |
| Guild | 3 | 43 | 15.08 | <0.0001 |

*Average Comparisons (Guild)*

Method of comparing pairs of means: Fisher. alpha=0.05

Mean Standard Difference Error = 0.274

Minimum average significant difference = 0.552

| **Coverage** | **Variable** | **Guild** | **Stocking** | **USA** | **N** | **Group** |
| --- | --- | --- | --- | --- | --- | --- |
| Semi-open | FDis | INS | 3.25 | 0.18 | 13 | A |
| Semi-open | FDis | GRA | 2.51 | 0.22 | 9 | B |
| Semi-open | FDis | FRU | 2.33 | 0.18 | 13 | B |
| Semi-open | FDis | SEE | 1.50 | 0.19 | 12 | C |

Stockings with a common letter are not significantly different (e > 0.05).

***Variable = RaoQ***

| **Fountain** | **GLNum** | **GLDen** | **F** | **P-Value** |
| --- | --- | --- | --- | --- |
| (Intercept) | 1 | 43 | 193.04 | <0.0001 |
| Guild | 3 | 43 | 16.54 | <0.0001 |

*Average Comparisons (Guild)*

Method of comparing pairs of means: Fisher. alpha=0.05

Mean Difference Mean Standard Error=1.61

Minimum average significant difference = 3.25

| **Coverage** | **Variable** | **Guild** | **Stocking** | **USA** | **N** | **Group** |
| --- | --- | --- | --- | --- | --- | --- |
| Semi-open | RaoQ | INS | 14.05 | 1.07 | 13 | A |
| Semi-open | RaoQ | GRA | 7.10 | 1.29 | 9 | B |
| Semi-open | RaoQ | FRU | 7.01 | 1.07 | 13 | B |
| Semi-open | RaoQ | SEE | 3.49 | 1.11 | 12 | C |

Stockings with a common letter are not significantly different (e > 0.05).

**Mosaics semi-closed vegetation (SC)**

***Variable = Richness***

| **Fountain** | **GLNum** | **GLDen** | **F** | **P-Value** |
| --- | --- | --- | --- | --- |
| (Intercept) | 1 | 95 | 170.81 | <0.0001 |
| Guild | 3 | 95 | 17.32 | <0.0001 |

*Average Comparisons (Guild)*

Method of comparing pairs of means: Fisher. alpha=0.05

Mean Difference Mean Standard Error=2.11

Average Minimum Significant Difference = 4.18

| **Coverage** | **Variable** | **Guild** | **Stocking** | **USA** | **N** | **Group** |
| --- | --- | --- | --- | --- | --- | --- |
| Semi-closed | Richness | INS | 18.29 | 1.38 | 28 | A |
| Semi-closed | Richness | FRU | 9.66 | 1.36 | 29 | B |
| Semi-closed | Richness | SEE | 6.24 | 1.60 | 21 | BC |
| Semi-closed | Richness | GRA | 4.76 | 1.60 | 21 | C |

Stockings with a common letter are not significantly different (e > 0.05).

***Variable = Abundance***

| **Fountain** | **GLNum** | **GLDen** | **F** | **P-Value** |
| --- | --- | --- | --- | --- |
| (Intercept) | 1 | 95 | 46.87 | <0.0001 |
| Guild | 3 | 95 | 4.50 | 0.0054 |

*Average Comparisons (Guild)*

Method of comparing pairs of means: Fisher. alpha=0.05

Mean difference average standard error=16.2

Minimum average significant difference = 32.2

| **Coverage** | **Variable** | **Guild** | **Stocking** | **USA** | **N** | **Group** |
| --- | --- | --- | --- | --- | --- | --- |
| Semi-closed | Individuals | INS | 66.43 | 10.66 | 28 | A |
| Semi-closed | Individuals | FRU | 53.00 | 10.48 | 29 | AB |
| Semi-closed | Individuals | SEE | 21.19 | 12.31 | 21 | BC |
| Semi-closed | Individuals | GRA | 16.48 | 12.31 | 21 | C |

Stockings with a common letter are not significantly different (e > 0.05).

***Variable = Dominance_D***

| **Fountain** | **GLNum** | **GLDen** | **F** | **P-Value** |
| --- | --- | --- | --- | --- |
| (Intercept) | 1 | 95 | 315.06 | <0.0001 |
| Guild | 3 | 95 | 16.16 | <0.0001 |

*Average Comparisons (Guild)*

Method of comparing pairs of means: Fisher. alpha=0.05

Mean difference average standard error=0.03

Minimum Average Significant Difference = 0.059

| **Coverage** | **Variable** | **Guild** | **Stocking** | **USA** | **N** | **Group** |
| --- | --- | --- | --- | --- | --- | --- |
| Semi-closed | Dominance_D | SEE | 0.29 | 0.02 | 21 | A |
| Semi-closed | Dominance_D | GRA | 0.19 | 0.02 | 21 | B |
| Semi-closed | Dominance_D | FRU | 0.17 | 0.02 | 29 | B |
| Semi-closed | Dominance_D | INS | 0.09 | 0.02 | 28 | C |

Stockings with a common letter are not significantly different (e > 0.05).

***Variable = Shannon_H***

| **Fountain** | **GLNum** | **GLDen** | **F** | **P-Value** |
| --- | --- | --- | --- | --- |
| (Intercept) | 1 | 95 | 1934.11 | <0.0001 |
| Guild | 3 | 95 | 39.38 | <0.0001 |

*Average Comparisons (Guild)*

Method of comparing pairs of means: Fisher. alpha=0.05

Mean difference average standard error=0.124

Minimum average significant difference = 0.245

| **Coverage** | **Variable** | **Guild** | **Stocking** | **USA** | **N** | **Group** |
| --- | --- | --- | --- | --- | --- | --- |
| Semi-closed | Shannon_H | INS | 2.67 | 0.08 | 28 | A |
| Semi-closed | Shannon_H | FRU | 1.97 | 0.08 | 29 | B |
| Semi-closed | Shannon_H | GRA | 1.53 | 0.09 | 21 | C |
| Semi-closed | Shannon_H | SEE | 1.52 | 0.09 | 21 | C |

Stockings with a common letter are not significantly different (e > 0.05).

***Variable = Margalef***

| **Fountain** | **GLNum** | **GLDen** | **F** | **P-Value** |
| --- | --- | --- | --- | --- |
| (Intercept) | 1 | 95 | 437.71 | <0.0001 |
| Guild | 3 | 95 | 29.47 | <0.0001 |

*Average Comparisons (Guild)*

Method of comparing pairs of means: Fisher. alpha=0.05

Mean Standard Difference Error = 0.33

Minimum Significant Difference Average = 0.655

| **Coverage** | **Variable** | **Guild** | **Stocking** | **USA** | **N** | **Group** |
| --- | --- | --- | --- | --- | --- | --- |
| Semi-closed | Margalef | INS | 4.22 | 0.22 | 28 | A |
| Semi-closed | Margalef | FRU | 2.28 | 0.21 | 29 | B |
| Semi-closed | Margalef | SEE | 1.76 | 0.25 | 21 | BC |
| Semi-closed | Margalef | GRA | 1.50 | 0.25 | 21 | C |

Stockings with a common letter are not significantly different (e > 0.05).

***Variable = FRic***

| **Fountain** | **GLNum** | **GLDen** | **F** | **P-Value** |
| --- | --- | --- | --- | --- |
| (Intercept) | 1 | 95 | 158.57 | <0.0001 |
| Guild | 3 | 95 | 4.40 | 0.0061 |

*Average Comparisons (Guild)*

Method of comparing pairs of means: Fisher. alpha=0.05

Mean difference average standard error=0.05

Minimum Average Significant Difference = 0.099

| **Coverage** | **Variable** | **Guild** | **Stocking** | **USA** | **N** | **Group** |
| --- | --- | --- | --- | --- | --- | --- |
| Semi-closed | FRic | FRU | 0.29 | 0.03 | 29 | A |
| Semi-closed | FRic | INS | 0.28 | 0.03 | 28 | A |
| Semi-closed | FRic | GRA | 0.16 | 0.04 | 21 | B |
| Semi-closed | FRic | SEE | 0.15 | 0.04 | 21 | B |

Stockings with a common letter are not significantly different (e > 0.05).

***Variable = FEve***

| **Fountain** | **GLNum** | **GLDen** | **F** | **P-Value** |
| --- | --- | --- | --- | --- |
| (Intercept) | 1 | 95 | 2809.57 | <0.0001 |
| Guild | 3 | 95 | 3.78 | 0.0131 |

*Average Comparisons (Guild)*

Method of comparing pairs of means: Fisher. alpha=0.05

Mean difference average standard error=0.036

Minimum Average Significant Difference = 0.072

| **Coverage** | **Variable** | **Guild** | **Stocking** | **USA** | **N** | **Group** |
| --- | --- | --- | --- | --- | --- | --- |
| Semi-closed | FAITH | INS | 0.72 | 0.02 | 28 | A |
| Semi-closed | FAITH | GRA | 0.70 | 0.03 | 21 | A |
| Semi-closed | FAITH | SEE | 0.70 | 0.03 | 21 | A |
| Semi-closed | FAITH | FRU | 0.62 | 0.02 | 29 | B |

Stockings with a common letter are not significantly different (e > 0.05).

***Variable = FDiv***

| **Fountain** | **GLNum** | **GLDen** | **F** | **P-Value** |
| --- | --- | --- | --- | --- |
| (Intercept) | 1 | 95 | 4905.08 | <0.0001 |
| Guild | 3 | 95 | 14.85 | <0.0001 |

*Average Comparisons (Guild)*

Method of comparing pairs of means: Fisher. alpha=0.05

Mean difference average standard error=0.03

Minimum Average Significant Difference = 0.059

| **Coverage** | **Variable** | **Guild** | **Stocking** | **USA** | **N** | **Group** |
| --- | --- | --- | --- | --- | --- | --- |
| Semi-closed | FDiv | GRA | 0.81 | 0.02 | 21 | A |
| Semi-closed | FDiv | FRU | 0.80 | 0.02 | 29 | A |
| Semi-closed | FDiv | SEE | 0.69 | 0.02 | 21 | B |
| Semi-closed | FDiv | INS | 0.65 | 0.02 | 28 | B |

Stockings with a common letter are not significantly different (e > 0.05).

***Variable = FDis***

| **Fountain** | **GLNum** | **GLDen** | **F** | **P-Value** |
| --- | --- | --- | --- | --- |
| (Intercept) | 1 | 95 | 932.15 | <0.0001 |
| Guild | 3 | 95 | 15.84 | <0.0001 |

*Average Comparisons (Guild)*

Method of comparing pairs of means: Fisher. alpha=0.05

Mean Difference Mean Standard Error=0.218

Minimum average significant difference = 0.432

| **Coverage** | **Variable** | **Guild** | **Stocking** | **USA** | **N** | **Group** |
| --- | --- | --- | --- | --- | --- | --- |
| Semi-closed | FDis | INS | 3.02 | 0.14 | 28 | A |
| Semi-closed | FDis | FRU | 2.52 | 0.14 | 29 | B |
| Semi-closed | FDis | GRA | 2.34 | 0.17 | 21 | B |
| Semi-closed | FDis | SEE | 1.53 | 0.17 | 21 | C |

Stockings with a common letter are not significantly different (e > 0.05).

***Variable = RaoQ***

| **Fountain** | **GLNum** | **GLDen** | **F** | **P-Value** |
| --- | --- | --- | --- | --- |
| (Intercept) | 1 | 95 | 228.66 | <0.0001 |
| Guild | 3 | 95 | 14.38 | <0.0001 |

*Average Comparisons (Guild)*

Method of comparing pairs of means: Fisher. alpha=0.05

Mean Difference Mean Standard Error=1.42

Minimum Average Significant Difference = 2.81

| **Coverage** | **Variable** | **Guild** | **Stocking** | **USA** | **N** | **Group** |
| --- | --- | --- | --- | --- | --- | --- |
| Semi-closed | RaoQ | INS | 12.42 | 0.93 | 28 | A |
| Semi-closed | RaoQ | FRU | 8.16 | 0.92 | 29 | B |
| Semi-closed | RaoQ | GRA | 6.29 | 1.08 | 21 | BC |
| Semi-closed | RaoQ | SEE | 3.43 | 1.08 | 21 | C |

Stockings with a common letter are not significantly different (e > 0.05).

**Mosaics Closed Vegetation (CL)**

***Variable = Richness***

| **Fountain** | **GLNum** | **GLDen** | **F** | **P-Value** |
| --- | --- | --- | --- | --- |
| (Intercept) | 1 | 51 | 269.54 | <0.0001 |
| Guild | 3 | 51 | 24.73 | <0.0001 |

*Average Comparisons (Guild)*

Method of comparing pairs of means: Fisher. alpha=0.05

Mean Difference Mean Standard Error=1.61

Minimum Average Significant Difference = 3.24

| **Coverage** | **Variable** | **Guild** | **Stocking** | **USA** | **N** | **Group** |
| --- | --- | --- | --- | --- | --- | --- |
| Closed | Richness | INS | 16.81 | 1.04 | 16 | A |
| Closed | Richness | FRU | 10.38 | 1.04 | 16 | B |
| Closed | Richness | GRA | 5.36 | 1.26 | 11 | C |
| Closed | Richness | SEE | 4.92 | 1.20 | 12 | C |

Stockings with a common letter are not significantly different (e > 0.05).

***Variable = Abundance***

| **Fountain** | **GLNum** | **GLDen** | **F** | **P-Value** |
| --- | --- | --- | --- | --- |
| (Intercept) | 1 | 51 | 48.62 | <0.0001 |
| Guild | 3 | 51 | 3.25 | 0.0291 |

*Average Comparisons (Guild)*

Method of comparing pairs of means: Fisher. alpha=0.05

Mean difference average standard error=16.2

Minimum average significant difference = 32.5

| **Coverage** | **Variable** | **Guild** | **Stocking** | **USA** | **N** | **Group** |
| --- | --- | --- | --- | --- | --- | --- |
| Closed | Abundance | INS | 59.00 | 10.47 | 16 | A |
| Closed | Abundance | FRU | 55.69 | 10.47 | 16 | A |
| Closed | Abundance | SEE | 29.17 | 12.08 | 12 | AB |
| Closed | Abundance | GRA | 15.82 | 12.62 | 11 | B |

Stockings with a common letter are not significantly different (e > 0.05).

***Variable = Dominance_D***

| **Fountain** | **GLNum** | **GLDen** | **F** | **P-Value** |
| --- | --- | --- | --- | --- |
| (Intercept) | 1 | 51 | 164.38 | <0.0001 |
| Guild | 3 | 51 | 21.44 | <0.0001 |

*Average Comparisons (Guild)*

Method of comparing pairs of means: Fisher. alpha=0.05

Mean difference average standard error=0.045

Minimum Average Significant Difference = 0.091

| **Coverage** | **Variable** | **Guild** | **Stocking** | **USA** | **N** | **Group** |
| --- | --- | --- | --- | --- | --- | --- |
| Closed | Dominance_D | SEE | 0.42 | 0.03 | 12 | A |
| Closed | Dominance_D | GRA | 0.19 | 0.04 | 11 | B |
| Closed | Dominance_D | FRU | 0.13 | 0.03 | 16 | BC |
| Closed | Dominance_D | INS | 0.08 | 0.03 | 16 | C |

Stockings with a common letter are not significantly different (e > 0.05).

***Variable = Shannon_H***

| **Fountain** | **GLNum** | **GLDen** | **F** | **P-Value** |
| --- | --- | --- | --- | --- |
| (Intercept) | 1 | 51 | 1675.37 | <0.0001 |
| Guild | 3 | 51 | 50.39 | <0.0001 |

*Average Comparisons (Guild)*

Method of comparing pairs of means: Fisher. alpha=0.05

Mean Standard Difference Error = 0.132

Minimum Significant Difference Average = 0.265

| **Coverage** | **Variable** | **Guild** | **Stocking** | **USA** | **N** | **Group** |
| --- | --- | --- | --- | --- | --- | --- |
| Closed | Shannon_H | INS | 2.67 | 0.09 | 16 | A |
| Closed | Shannon_H | FRU | 2.17 | 0.09 | 16 | B |
| Closed | Shannon_H | GRA | 1.65 | 0.10 | 11 | C |
| Closed | Shannon_H | SEE | 1.15 | 0.10 | 12 | D |

Stockings with a common letter are not significantly different (e > 0.05).

***Variable = Margalef***

| **Fountain** | **GLNum** | **GLDen** | **F** | **P-Value** |
| --- | --- | --- | --- | --- |
| (Intercept) | 1 | 51 | 545.72 | <0.0001 |
| Guild | 3 | 51 | 36.57 | <0.0001 |

*Average Comparisons (Guild)*

Method of comparing pairs of means: Fisher. alpha=0.05

Mean difference average standard error=0.289

Minimum average significant difference = 0.581

| **Coverage** | **Variable** | **Guild** | **Stocking** | **USA** | **N** | **Group** |
| --- | --- | --- | --- | --- | --- | --- |
| Closed | Margalef | INS | 4.05 | 0.19 | 16 | A |
| Closed | Margalef | FRU | 2.50 | 0.19 | 16 | B |
| Closed | Margalef | GRA | 1.63 | 0.23 | 11 | C |
| Closed | Margalef | SEE | 1.39 | 0.22 | 12 | C |

Stockings with a common letter are not significantly different (e > 0.05).

***Variable = FRic***

| **Fountain** | **GLNum** | **GLDen** | **F** | **P-Value** |
| --- | --- | --- | --- | --- |
| (Intercept) | 1 | 51 | 122.36 | <0.0001 |
| Guild | 3 | 51 | 4.58 | 0.0065 |

*Average Comparisons (Guild)*

Method of comparing pairs of means: Fisher. alpha=0.05

Mean difference average standard error=0.054

Minimum Average Significant Difference = 0.109

| **Coverage** | **Variable** | **Guild** | **Stocking** | **USA** | **N** | **Group** |
| --- | --- | --- | --- | --- | --- | --- |
| Closed | FRic | INS | 0.30 | 0.04 | 16 | A |
| Closed | FRic | FRU | 0.26 | 0.04 | 16 | A |
| Closed | FRic | GRA | 0.19 | 0.04 | 11 | AB |
| Closed | FRic | SEE | 0.11 | 0.04 | 12 | B |

Stockings with a common letter are not significantly different (e > 0.05).

***Variable = FEve***

| **Fountain** | **GLNum** | **GLDen** | **F** | **P-Value** |
| --- | --- | --- | --- | --- |
| (Intercept) | 1 | 51 | 1322.41 | <0.0001 |
| Guild | 3 | 51 | 5.50 | 0.0024 |

*Average Comparisons (Guild)*

Method of comparing pairs of means: Fisher. alpha=0.05

Mean difference average standard error=0.051

Minimum average significant difference = 0.102

| **Coverage** | **Variable** | **Guild** | **Stocking** | **USA** | **N** | **Group** |
| --- | --- | --- | --- | --- | --- | --- |
| Closed | FAITH | GRA | 0.71 | 0.04 | 11 | A |
| Closed | FAITH | INS | 0.69 | 0.03 | 16 | A |
| Closed | FAITH | SEE | 0.66 | 0.04 | 12 | A |
| Closed | FAITH | FRU | 0.53 | 0.03 | 16 | B |

Stockings with a common letter are not significantly different (e > 0.05).

***Variable = FDiv***

| **Fountain** | **GLNum** | **GLDen** | **F** | **P-Value** |
| --- | --- | --- | --- | --- |
| (Intercept) | 1 | 51 | 1800.60 | <0.0001 |
| Guild | 3 | 51 | 4.71 | 0.0056 |

*Average Comparisons (Guild)*

Method of comparing pairs of means: Fisher. alpha=0.05

Mean Standard Difference Error = 0.047

Minimum Average Significant Difference = 0.095

| **Coverage** | **Variable** | **Guild** | **Stocking** | **USA** | **N** | **Group** |
| --- | --- | --- | --- | --- | --- | --- |
| Closed | FDiv | FRU | 0.79 | 0.03 | 16 | A |
| Closed | FDiv | GRA | 0.75 | 0.04 | 11 | AB |
| Closed | FDiv | INS | 0.67 | 0.03 | 16 | BC |
| Closed | FDiv | SEE | 0.64 | 0.04 | 12 | C |

Stockings with a common letter are not significantly different (e > 0.05).

***Variable = FDis***

| **Fountain** | **GLNum** | **GLDen** | **F** | **P-Value** |
| --- | --- | --- | --- | --- |
| (Intercept) | 1 | 51 | 1488.13 | <0.0001 |
| Guild | 3 | 51 | 51.44 | <0.0001 |

*Average Comparisons (Guild)*

Method of comparing pairs of means: Fisher. alpha=0.05

Mean Standard Difference Average Error=0.153

Minimum average significant difference = 0.307

| **Coverage** | **Variable** | **Guild** | **Stocking** | **USA** | **N** | **Group** |
| --- | --- | --- | --- | --- | --- | --- |
| Closed | FDis | INS | 2.94 | 0.10 | 16 | A |
| Closed | FDis | GRA | 2.23 | 0.12 | 11 | B |
| Closed | FDis | FRU | 2.10 | 0.10 | 16 | B |
| Closed | FDis | SEE | 1.07 | 0.11 | 12 | C |

Stockings with a common letter are not significantly different (e > 0.05).

***Variable = RaoQ***

| **Fountain** | **GLNum** | **GLDen** | **F** | **P-Value** |
| --- | --- | --- | --- | --- |
| (Intercept) | 1 | 51 | 368.08 | <0.0001 |
| Guild | 3 | 51 | 38.45 | <0.0001 |

*Average Comparisons (Guild)*

Method of comparing pairs of means: Fisher. alpha=0.05

Mean Standard Difference Error = 0.916

Minimum average significant difference = 1.84

| **Coverage** | **Variable** | **Guild** | **Stocking** | **USA** | **N** | **Group** |
| --- | --- | --- | --- | --- | --- | --- |
| Closed | RaoQ | INS | 11.35 | 0.59 | 16 | A |
| Closed | RaoQ | GRA | 5.91 | 0.71 | 11 | B |
| Closed | RaoQ | FRU | 5.74 | 0.59 | 16 | B |
| Closed | RaoQ | SEE | 1.86 | 0.68 | 12 | C |

Stockings with a common letter are not significantly different (e > 0.05).

References and citations of the R packages on which this module is based

{nlme}: Linear and Nonlinear Mixed Effects Models Jose Pinheiro Douglas Bates Saikat DebRoy Deepayan Sarkar R Core Team 2021 R package version 3.1-152 <https://CRAN.R-project.org/package=nlme>

Fisher. R.A. (1935). The Design of Experiments. Oliver & Boyd. Edinburgh.

Navure Team (2023). Navure (2.8.2): A data-science-statistic oriented application for making evidence-based decisions. URL <http://www.navure.com>

Pinheiro. J.. & Bates. D. (2006). Mixed-effects models in S and S-PLUS. Springer science & business media.

R 3.6.3: A Language and Environment for Statistical Computing R Core Team R Foundation for Statistical Computing Vienna. Austria 2020 <https://www.R-project.org/>
